# Structural and molecular basis of choline uptake into the brain by FLVCR2

**DOI:** 10.1101/2023.10.05.561059

**Authors:** Rosemary J. Cater, Dibyanti Mukherjee, Eva Gil Iturbe, Satchal K. Erramilli, Ting Chen, Katie Koo, Nicolás Santander Grez, Andrew Reckers, Brian Kloss, Tomasz Gawda, Brendon C. Choy, Zhening Zheng, Oliver B. Clarke, Sook Wah Yee, Anthony A. Kossiakoff, Matthias Quick, Thomas Arnold, Filippo Mancia

## Abstract

Choline is an essential nutrient that the human body needs in vast quantities for cell membrane synthesis, epigenetic modification, and neurotransmission. The brain has a particularly high demand for choline, but how it enters the brain has eluded the field for over fifty years. The MFS transporter FLVCR1 was recently determined to be a choline transporter, and while this protein is not highly expressed at the blood-brain barrier (BBB), its relative FLVCR2 is. Previous studies have shown that mutations in human *Flvcr2* cause cerebral vascular abnormalities, hydrocephalus, and embryonic lethality, but the physiological role of FLVCR2 is unknown. Here, we demonstrate both *in vivo* and *in vitro* that FLVCR2 is a BBB choline transporter and is responsible for the majority of choline uptake into the brain. We also determine the structures of choline-bound FLVCR2 in the inward- and outward-facing states using cryo-electron microscopy to 2.49 and 2.77 Å resolution, respectively. These results reveal how the brain obtains choline and provide molecular-level insights into how FLVCR2 binds choline in an aromatic cage and mediates its uptake. Our work could provide a novel framework for the targeted delivery of neurotherapeutics into the brain.

## Main

Choline is an essential nutrient that serves various metabolic roles that are critical for healthy brain development and function^1–3^. It is a precursor to phosphatidylcholine, which is required in enormous quantities for membrane synthesis in all cells^4,5^, as well as betaine, a critical metabolite involved in the methylation of DNA and protein^5,6^. In cholinergic neurons, it is also converted to acetylcholine, a neurotransmitter driving many central and peripheral nervous system functions including memory and muscle contraction^5^. Despite the brain’s high demand for choline, it cannot be efficiently synthesised *de novo*^3^. Instead, the brain derives the majority of its choline from systemic circulation, most of which originates from dietary sources^1,3^. Indeed, *in vivo* radioactive tracer studies have shown that choline is transported rapidly and specifically across the blood-brain barrier (BBB) without metabolic alteration by a saturable carrier-mediated process^7^.

The identity of a BBB choline transporter and its associated mechanism have eluded the field for 50 years. Several candidate transporters have been considered, including members of the SLC22/OCT and SLC44/CTL families. However, these proteins exhibit low affinity for choline and/or are not highly expressed at the BBB^8–11^ (**Extended Data Fig. 1a**). The only two *bona fide* human choline transporters identified thus far are the Major Facilitator Superfamily (MFS) transporter FLVCR1 (also known as *MFSD7B* or *SLC49A1*), which is expressed in most cell types but not highly enriched in brain endothelial cells^12,13^, and the high-affinity choline transporter ChT (*SLC5A7*), which is almost exclusively expressed in cholinergic neurons^14^ (**Extended Data Fig. 1a**).

FLVCR2 (also known as *MFSD7C* or *SLC49A2*) is a close relative of FLVCR1 (55% identity) that is highly and specifically expressed in BBB endothelial cells throughout development and into adulthood^15,16^ (**Extended Data Fig. 1a**). In humans, mutations in FLVCR2 cause proliferative vasculopathy and hydranencephaly hydrocephalus (PVHH; OMIM 225790), also known as Fowler syndrome. This rare autosomal recessive brain vascular disorder is associated with impaired cerebral angiogenesis, hydrocephalus, and embryonic lethality^17^. Endothelial cell-specific knockout of *Flvcr2* in mice recapitulates human PVHH phenotypes, demonstrating that endothelial-expressed FLVCR2 is critical for both cerebral angiogenesis and normal brain development^16,18^. However, beyond this genetic association, the physiological role of FLVCR2 at the BBB, as well as the identity of its substrate, are unknown.

Here, we show that FLVCR2 is a BBB choline transporter, and present substrate-bound structures of this protein in both the outward-facing state (OFS; 2.49 Å resolution) and inward-facing state (IFS; 2.75 Å resolution), which we determined using single-particle cryo-electron microscopy (cryo-EM). These findings de-orphan FLVCR2 and provide molecular insights into its mechanism of choline transport.

## FLVCR2 is a blood-brain barrier choline transporter

Our previous mRNA expression profiling experiments in mice showed that *Flvcr2* is strongly and specifically expressed in BBB endothelial cells during development and into adulthood^16^. To confirm FLVCR2 expression at the protein level, and determine its subcellular localisation *in vivo*, we attached a hemagglutinin (HA) tag to the C-terminus of FLVCR2 in mice using CRISPR-Cas9 (*Flvcr2^HA/HA^* mice). Immunolocalisation in adult *Flvcr2^HA/HA^* mice indicated that FLVCR2 is exclusively found in endothelial cells of all brain vascular segments (artery/capillary/vein; **Fig. 1a, left panel**). Furthermore, FLVCR2 predominantly co-localised with intercellular adhesion molecule 2 (ICAM2) rather than with collagen type IV (COLIV), indicating that it is more highly expressed on the luminal (blood-facing) membrane than the abluminal (brain-facing) membrane (**Fig. 1a, right panel**). This luminal expression places FLVCR2 in an ideal location for the uptake of choline from the circulation.

**Figure 1.**
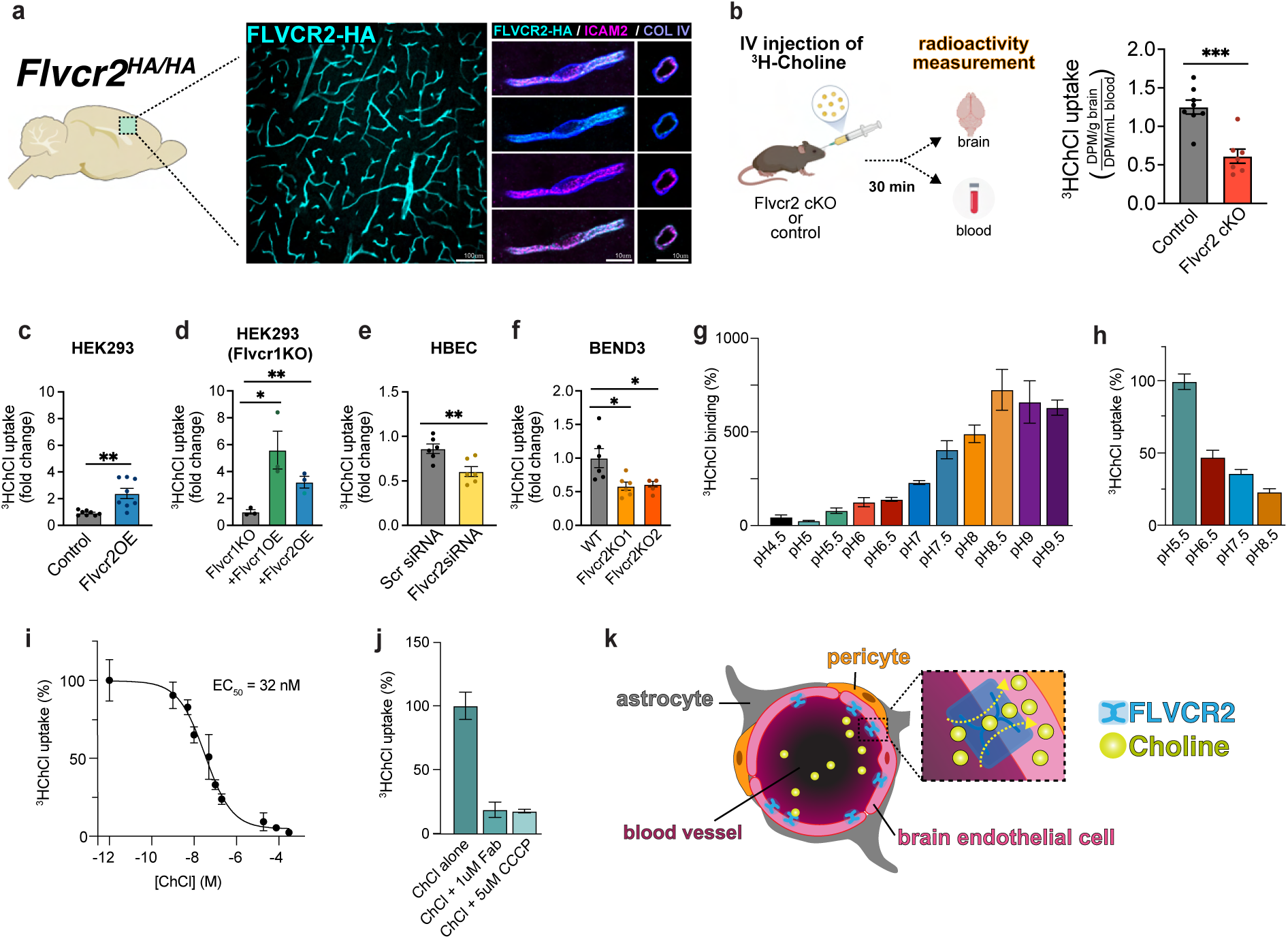
FLVCR2 is a blood-brain barrier choline transporter. **a,** Fixed sections from *Flvcr2*^HA/HA^ mice were stained for HA (cyan) and pan-endothelial markers ICAM2 (pink) and COL IV (purple), which mark the luminal (blood) and abluminal (brain) sides of the blood vessel. The rightmost panel depicts side-on and frontal views of the vascular segments at high resolution. Scale bars are shown. **b,** Control (*Flvcr2^Fl/+^ Cdh5Cre^ER^*) and *Flvcr2* conditional KO (cKO; *Flvcr2^Fl/Fl^ Cdh5Cre^ER^*) mice were injected with the same dose of [^3^H]ChCl intravenously. Blood and brains were collected 30 min after injection and d.p.m. was quantified by scintillation. Uptake is calculated as d.p.m. per g brain/d.p.m. per mL serum and then normalised to control. Data are mean ± s.e.m. (n = 12) combined from two independent experiments; **** indicates p < 0.0001. **c,** [^3^H]ChCl uptake in un-transfected HEK293 cells (control) and HEK293 cells transfected with *Mm*FLVCR2 cDNA (*Flvcr2* OE; n = 11); ** indicates p <0.01. **d,** [^3^H]ChCl uptake in *Flvcr1* KO, *Flvcr1* KO + *Flvcr1* OE and *Flvcr1* KO + *Flvcr2* OE cells (n = 4); *** indicates p <0.001 and ** indicates p < 0.01. **e,** Human brain microvascular endothelial cells (HBECs) transfected with either scr siRNA or *Flvcr2* siRNA (n = 10); **** indicates p < 0.000. **f**, BEND3.1 WT and FLVCR2 KO cells (KO1, KO2; n = 5); ** indicates p < 0.01 and *** indicates p <0.001. Uptake is normalised as a proportion of WT mean ± s.e.m. **g,** pH-dependence of 1 µM [^3^H]ChCl binding to detergent-purified *Mm*FLVCR2. Binding was performed with the scintillation proximity assay^55^. Data represent the mean ± s.e.m. (n = 3) specific signal (total c.p.m. minus c.p.m. in the presence of 800 mM imidazole) and are normalised to the specific signal measured at pH 5.5. **h,** Uptake of 1 µM [^3^H]ChCl into *Mm*FLVCR2-containing proteoliposomes after 1 minute. The external pH was varied as indicated while maintaining an internal pH of 7.5. Data are mean ± s.e.m. (n = 3). For all liposomal uptake assays the specific uptake activity of *Mm*FLVCR2 was determined as detailed in the methods. **i,** Uptake of 0.1 µM [^3^H]ChCl into *Mm*FLVCR2-containing proteoliposomes measured after 1 minute in the presence of increasing concentrations of unlabelled ChCl. The external and internal pH buffers were 5.5 and 7.5, respectively. Data yielded a log(EC_50_) = −7.48 ± 0.09 (corresponding to a mean of 32 nM). Data are mean ± s.e.m. (n = 4). **j,** Uptake of 1 µM [^3^H]ChCl into *Mm*FLVCR2-containing proteoliposomes after 1 minute measured in the presence or absence of either 1 μM Fab FLV23 or 5 μM CCCP. The external and internal pH buffers were 5.5 and 7.5, respectively. Data are mean ± s.e.m. (n = 3). **k,** Schematic representation of the role FLVCR2 plays in choline uptake at the BBB. FLVCR2 is expressed on the luminal membrane of brain microvascular endothelial cells where it mediates uptake of choline. Statistical analysis was performed using GraphPad Prism t-tests (two conditions) or ANOVA (three or more conditions).

To determine whether FLVCR2 transports choline across the BBB *in vivo*, we performed assays that measured the uptake of intravenous radiolabelled choline ([^3^H]ChCl) into the brains of mice that were genetically deficient in vascular endothelial *Flvcr2* (*Flvcr2^fl/fl^;Cdh5^CreER^*) and their littermate controls (*Flvcr2^fl/+^;Cdh5^CreER^*). After 30 minutes – the time required to reach saturation of choline in the brain^7^ – we found that radiolabelled choline accretion was significantly reduced in the *Flvcr2* conditional knockout (cKO) mice compared to the control group (**Fig. 1b**).

To further characterise FLVCR2-mediated choline transport, we established a number of cell-based uptake assays. First, we heterologously expressed *Flvcr2* in HEK293 cells. Uptake of [^3^H]ChCl after a 30-minute incubation was three times the uptake in non-transfected cells (**Fig. 1c**). As FLVCR1 is a recently characterised choline transporter^12^, we sought to compare choline uptake attributed to over-expression of *Flvcr1* versus *Flvcr2*. *Flvcr1* is endogenously expressed in HEK293 cells (**Extended Data Fig. 1b**), and we thus transiently transfected *Flvcr1* KO HEK293 cells with *Flvcr1* and *Flvcr2*. Over-expression of both *Flvcr1* and *Flvcr2* led to significant increases in choline uptake (**Fig. 1d**). In line with these results, siRNA-mediated knock-down of *Flvcr2* in immortalised human brain endothelial cells (HBEC) and CRISPR-Cas9-mediated knockout of *Flvcr2* from immortalised mouse brain endothelial (BEND3) cells caused significant reductions in [^3^H]ChCl uptake (34% and 25%, respectively; **Fig. 1e, f**). Taken together, these data demonstrate that FLVCR2 transports choline into BBB endothelial cells and is responsible for brain uptake of choline.

To establish FLVCR2 binding and transport kinetics, we purified FLVCR2 for use in binding and uptake experiments in a more isolated *in vitro* setting. To obtain purified protein, we screened six GFP-tagged FLVCR2 orthologs for expression and stability. We identified FLVCR2 from *Mus musculus* (*Mm*FLVCR2) – which shares 87% sequence identity with human FLVCR2 – as the most stable and well-expressing ortholog (**Extended Data Fig. 3a and 4**). Following optimisation of recombinant protein expression and purification (**Extended Data Fig. 3b**), we measured binding of [^3^H]ChCl to detergent-solubilised *Mm*FLVCR2 by scintillation proximity assays^19^. The results of these experiments demonstrate that *Mm*FLVCR2 can bind choline specifically, and that binding is pH-dependent, with the lowest levels of binding observed at pH 5.5, and the highest at pH 8.5 (**Fig. 1g**). To characterise choline transport kinetics, we then reconstituted *Mm*FLVCR2 into liposomes. We performed [^3^H]ChCl uptake assays in the presence of pH gradients that were either inwardly directed (external buffer pH 5.5/6.5; internal buffer pH 7.5) or outwardly directed (external buffer pH 8.5; internal buffer pH 7.5), and in the absence of any gradient (external and internal buffers pH 7.5). With an external buffer at pH 5.5, [^3^H]ChCl uptake levels were three times greater than those in the absence of a pH gradient, and four times greater than those in the presence of an outwardly-directed gradient (**Fig. 1h**). These data confirm that FLVCR2 transports choline, and suggest that this process may be proton-dependent, a feature commonly observed amongst MFS transporters^20^. Isotopic dilution assays yielded a mean half-maximal binding constant (EC_50_) of 32 nM for choline (**Fig. 1i**). Notably, the plasma concentration of free choline in human adult serum is ∼8-10 µM^21^, and thus this EC_50_ is sufficient to support the efficient uptake of choline from the circulation into brain endothelial cells. To assess the effect of pH on [^3^H]ChCl uptake by FLVCR2, we used the proton-specific ionophore CCCP to dissipate the inwardly-directed proton gradient across the liposomal membrane^22^. In the presence of CCCP, the efficiency of [^3^H]ChCl uptake was five times lower than uptake in its absence, and was similar to that observed in the presence of an antigen-binding fragment (Fab) which binds to a region of FLVCR2 that prevents structural rearrangements required for transport (**Fig. 1j, Extended Data Fig. 7b**).

Taken together, these results add FLVCR2 to the short list of *bona fide* human choline transporters and demonstrate that it is expressed in brain endothelial cells, where it mediates uptake of choline across the BBB into the brain (**Fig. 1k**).

## The structure of FLVCR2 in the outward-facing state

To understand FLVCR2-mediated choline transport at an atomic level, we employed cryo-EM to determine the structure of *Mm*FLVCR2 (hereafter referred to simply as FLVCR2) in complex with choline, its newly defined substrate. FLVCR2 is just 60 kDa in size and is not predicted to have substantial extra-membrane features, making it a difficult target for structure determination using cryo-EM^23^. To overcome this hurdle, we used a synthetic phage display library^24–26^ to identify FLVCR2-specific, high affinity Fabs (∼50 kDa in size) to generate a complex with FLVCR2. This approach aids cryo-EM structure determination by increasing the size of the imaged particles and facilitating particle alignment^27,28^. A total of 40 candidate Fabs were identified, including FLV23, our initial Fab of choice (**Extended Data Fig. 3c-e**). Cryo-EM data analysis of detergent-purified FLVCR2 in complex with 1 mM choline and Fab FLV23 yielded high-quality two-dimensional class averages with clear individual transmembrane helices (TMs; **Extended Data Fig. 5a**). These data ultimately allowed for reconstruction of a 2.49 Å resolution cryo-EM density map (**Fig. 2a, Extended Data Fig. 5** and **Extended Data Table 1**). We built a model of FLVCR2 into this map that includes residues 99-312 and 319-520 of the 551 in total (**Fig. 2b** and **Extended Data Fig. 6**). As anticipated, the 98-amino acid N-terminus and 31-amino acid C-terminus, which are predicted to be disordered, were not observed in our map.

**Figure 2.**
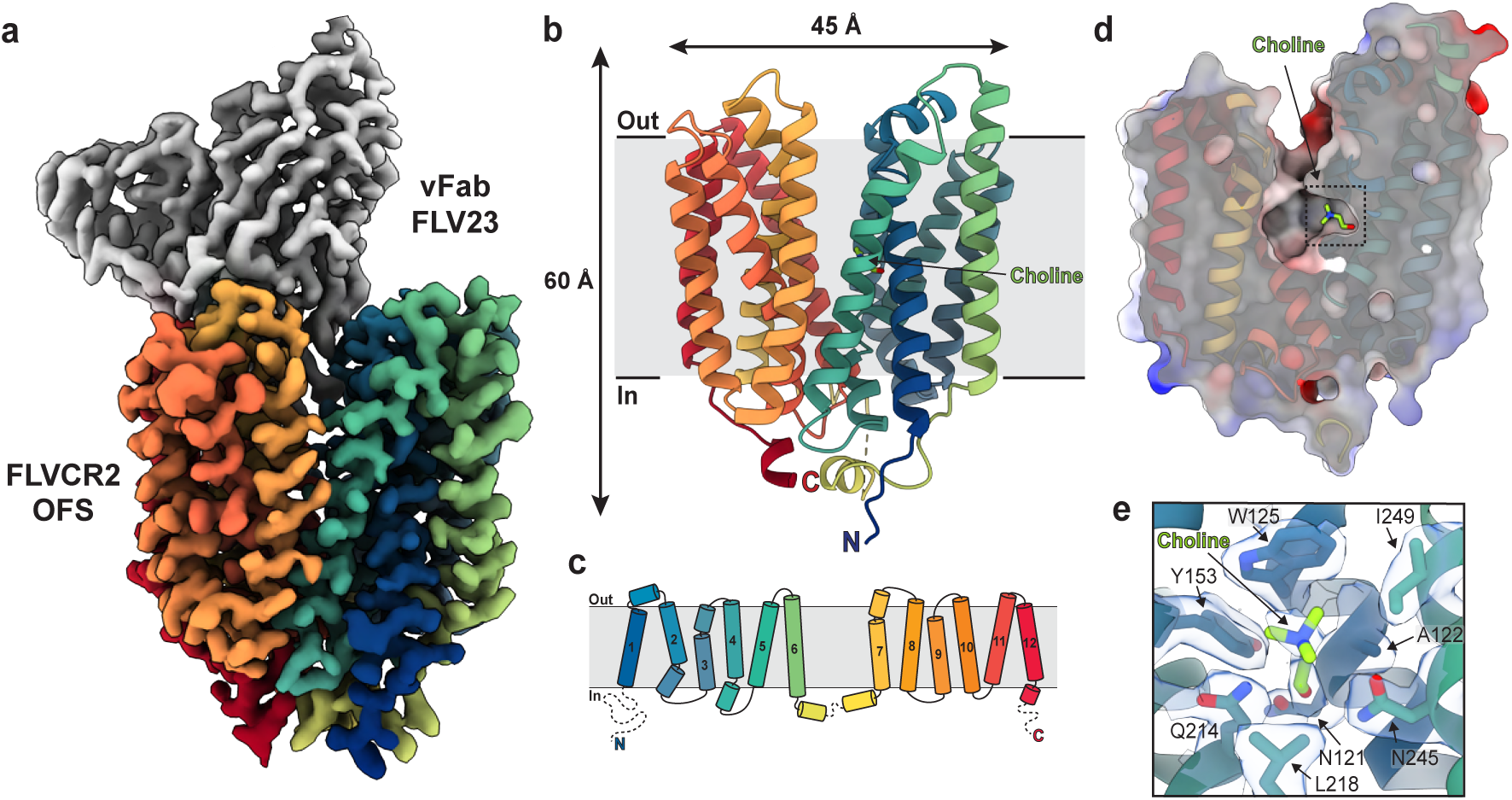
The structure of FLVCR2 in an outward-facing conformation. **a,** The 2.49 Å resolution cryo-EM density map of FLVCR2 in complex with Fab FLV23. Density corresponding to FLVCR2 is shown in rainbow from the N-terminus (blue) to the C-terminus (red) and in grey for the variable region of Fab FLV23. **b,** The OFS structure of FLVCR2 in the plane of the membrane, coloured as in **a**. Choline is shown in green stick representation and an unresolved intracellular loop as a dashed line. **c,** Topology of FLVCR2 showing the twelve numbered TM helices, coloured as in **a**. Disordered regions that could not be resolved in our cryo-EM map are shown as dashed lines. **d,** Surface representation of the electrostatic potential of the extracellular cavity. Red and blue indicate negatively and positively charged regions, respectively. Inset refers to the region highlighted in panel **e**. The structure is overlaid in rainbow ribbon and choline is shown in green stick representation. **e,** Choline binding site in the OFS. Choline and surrounding residues are shown in stick representation and the cryo-EM density corresponding to these is displayed as a semi-transparent light blue surface.

FLVCR2 has a classical MFS transporter fold comprising twelve TMs arranged into two pseudo-symmetric six-helix bundles (the N- and C-domains) that are connected by a flexible cytoplasmic loop^20^ (**Fig. 2b, c**). In this structure, the transporter is in the OFS, and exhibits a large extracellular cavity that extends approximately halfway across the membrane (**Fig. 2b, d, Extended Data Fig. 7a**). This cavity is lined primarily by aromatic, hydrophobic, and polar amino acids and notably lacks any positively or negatively charged residues (**Fig. 2d, Extended Data Fig. 7a**). We observed a non-protein cryo-EM density in a pocket within the extracellular cavity lined by N121, A122, W125, Y153, Q214, L218, N245, and I249 from the N-domain. Given that the equivalent residue of Y153 in FLVCR1 is critical for choline uptake^13^, we reasoned that this density likely corresponded to choline and fit it accordingly (**Fig. 2e**). Here, the trimethylammonium group of choline is coordinated by W125 and Y153 through cation-π interactions, while the hydroxyl group forms a hydrogen bond with N121 (**Fig. 2b, d, e**). Other residues surrounding choline at the site include A122, Q214, L218, N245, and I249. Notably, the C-domain is not involved in coordination of choline in this conformation.

## The structure of FLVCR2 in the inward-facing state

To further our understanding of FLVCR2-mediated choline transport, we next sought to obtain a structure of FLVCR2 in the IFS. As an MFS transporter, FLVCR2 likely utilises the rocker-switch mechanism to transport choline^20^. Its transition from the OFS to the IFS will therefore involve a conformational change which in part requires the pivoting of TM2 and TM11 towards each other at the membrane’s outer edge. In our OFS structure, Fab FLV23 is bound at this site (**Extended Data Fig. 7b**). This observation, together with our functional data showing that this Fab inhibits FLVCR2-mediated choline transport (**Fig. 1h**), suggests that this Fab impedes the conformational changes required to enter the IFS. We therefore screened for a Fab that binds to a different epitope to allow structure determination of the IFS. To this end, we performed epitope binning assays, which revealed that only one of our 40 FLVCR2-specific Fabs – Fab FLV9 – bound to an epitope distinct from that of FLV23 (**Extended Data Fig. 3f**). Structure determination of detergent-purified FLVCR2 in complex with 1 mM ChCl and FLV9 again led to high-quality two-dimensional class averages and ultimately yielded a density map with a resolution of 2.77 Å (**Fig. 3a, Extended Data Fig. 9 and Extended Data Table 2**). We built a model of FLVCR2 into this map comprising residues 100-312 and 319-520 of the 551 total (**Fig. 3** and **Extended Data Fig. 10**), which revealed FLVCR2 in the IFS (**Fig. 3b**). This inward-facing conformation features a large intracellular cavity. Similar to the extracellular cavity in the OFS, this cavity is lined almost entirely by non-charged residues – with the exception of R223, E447, E458, and R516, which are located on its intracellular edge (**Fig. 3c, Extended Data Fig. 7a**). At the deepest part of this cavity, a well-defined non-protein cryo-EM density was observed, into which we fit a choline molecule (**Fig. 3d**). Here, residues W125, Y153, and Y348 form an “aromatic cage” that surrounds the trimethylammonium group of the choline, and residues M154, Q214, and Q470 provide a hydrogen-bonding network to coordinate the hydroxyl end of the molecule (**Fig. 3d)**. This choline binding site bears clear resemblance to that of the recently determined *S. pneumoniae* choline transporter LicB – the only other choline transporter structure reported to date^29^. Given this similarity, and the ability of LicB to transport substrates other than choline (specifically, acetylcholine and arsenocholine), we decided to assess the promiscuity of FLVCR2’s substrate binding site. To achieve this, we performed a competition assay between [^3^H]ChCl and unlabelled choline-like metabolites including acetylcholine, betaine, and ethanolamine, as well as unlabelled choline itself. This experiment demonstrated that acetylcholine and betaine bind to FLVCR2 with a similar affinity to choline (**Fig. 3e**).

**Figure 3.**
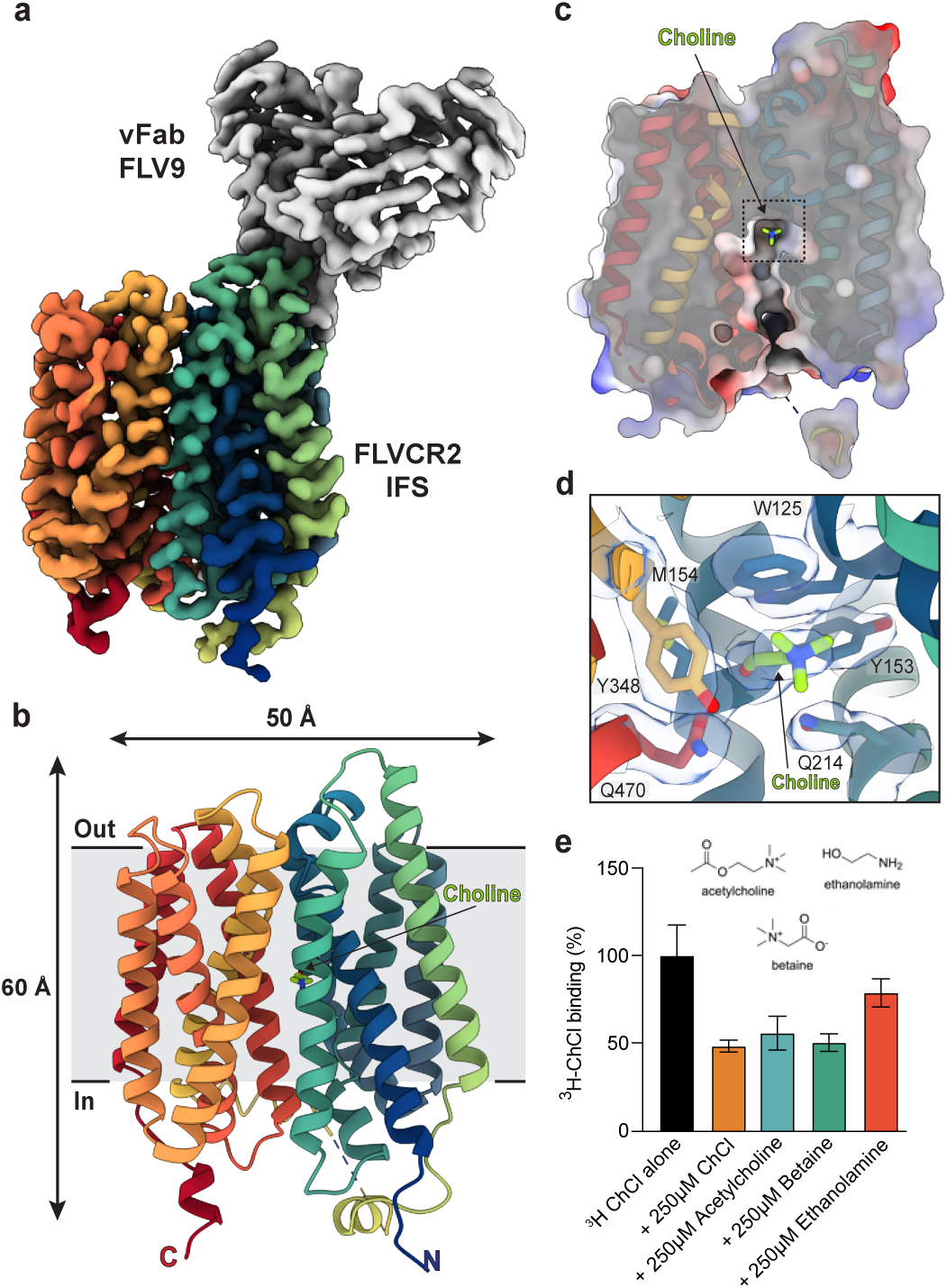
The structure of FLVCR2 in an inward-facing conformation. **a,** The 2.77 Å resolution cryo-EM density map of FLVCR2 in complex with Fab FLV9. Density corresponding to FLVCR2 is shown in rainbow from the N-terminus (blue) to the C-terminus (red) and in grey for the variable region of Fab FLV9. **b,** The IFS structure of FLVCR2 in the plane of the membrane, coloured as in **a**. Choline is shown in green stick representation and an unresolved intracellular loop as a dashed line. **c,** Surface representation of the electrostatic potential of the intracellular cavity. Red and blue indicate negatively and positively charged regions, respectively. Inset refers to the region highlighted in panel **d**. The structure is overlaid in rainbow ribbon and choline is shown in green stick representation. **d,** Choline binding site in the IFS. Choline and surrounding residues are shown in stick representation and the cryo-EM density corresponding to these is displayed as a semi-transparent light blue surface. **e,** Specific binding of 1 µM [^3^H]ChCl alone or in the presence of 250 µM ChCl, acetylcholine, betaine, or ethanolamine, with their chemical structures shown above. Data are mean ± s.e.m. (n = 3) of the specific signal (total c.p.m. minus the c.p.m. in the presence of 800 mM imidazole) and normalised to the specific signal in the absence of any unlabelled substrate.

## Discussion

We have determined that FLVCR2 is a BBB choline transporter, and have solved its structure in the OFS and IFS with choline bound in the central cavity (**Fig. 2, 3, 4a**). In the OFS, choline is bound exclusively by the N-domain, with coordination primarily provided by residues N121, W125, and Y153. Upon transition to the IFS, the trimethylammonium group of choline remains in the same location, but the hydroxyl group is reoriented as residues from the C-domain, including Y348 and Q470, move down to become available for substrate coordination (**Fig. 4b**). Notably, all residues involved in substrate coordination are highly conserved across mammalian FLVCR2 orthologs (**Extended Data Fig. 4, 7c**), as well as across mammalian FLVCR1 species, with the exception of I249, which is a threonine in FLVCR1 (**Extended Data Fig. 4**).

**Figure 4.**
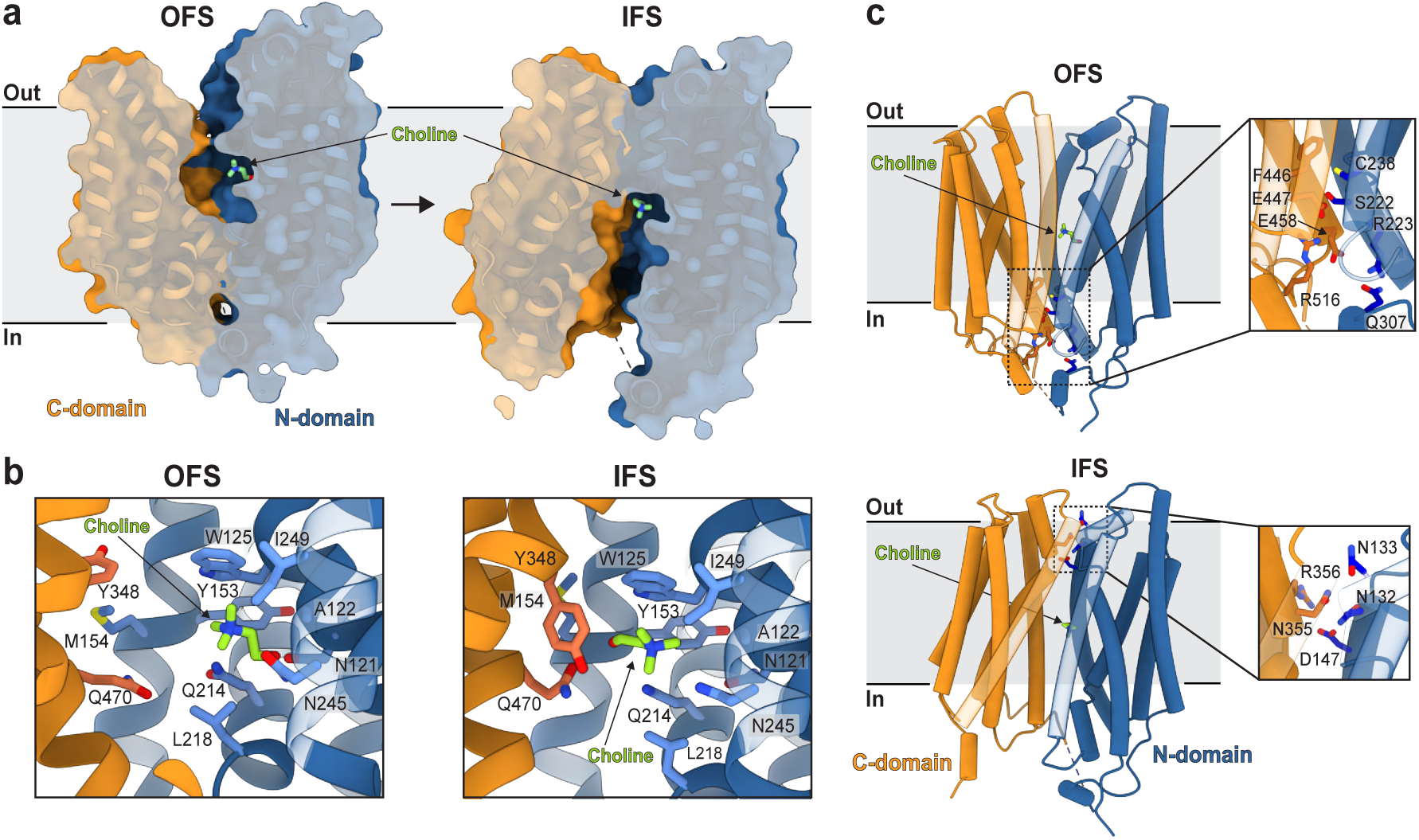
Mechanism of FLVCR2-mediated choline transport. **a,** Cross-sectional surface representation of FLVCR2 in the OFS (left) and IFS (right). The N- and C-domains are coloured in blue and orange, respectively. Choline is shown in green stick representation. **b,** Choline binding site in the OFS (left) and IFS (right). The protein is coloured as in panel **a** and shown in cartoon representation with residues involved in choline coordination from the N- and C-domain shown as blue and red-orange sticks, respectively. Choline is also shown in green stick representation. TMs in the foreground are shown in partial transparency for visual clarity. **c,** FLVCR2 in the OFS (top) and IFS (bottom) with residues participating in interdomain interactions highlighted in stick representation. The protein is coloured as in panel **b** with TMs shown as tubes and those in the foreground in partial transparency for visual clarity.

Analysis of the OFS and IFS structures reveals distinct networks of charged/polar residues that hold together the N- and C-domains in each state. In the OFS, this network comprises residues S222, R223, C238, and Q307 from the N-domain and E447, E458, and R516 from the C-domain. Residue F446 sits directly above this network, where it defines the base of the extracellular cavity (**Fig. 4c**, upper panel). In the IFS, inter-domain contacts are formed between N132, N133, and D147 from the N-domain and N355 and R356 from the C-domain (**Fig. 4c**, lower panel). Interestingly, human mutations of the residues corresponding to N132 and R356 (N109K and R333C, respectively; **Extended Data Fig. 4, 8**) are two of the mutations associated with PVHH. Our structures reveal the important role that these residues play in stabilising the IFS and thus explain how these mutations could disrupt choline transport and manifest in PVHH symptomology.

Our structures of FLVCR2 demonstrate that the choline binding site contains a number of aromatic residues. Aromatic interactions are a common mechanism for boosting substrate promiscuity at MFS binding sites^30,31^. In line with this, our results suggest that the FLVCR2 binding site is at least somewhat promiscuous (**Fig. 3e**). A better understanding of the full, detailed substrate profile of FLVCR2 (e.g., by screening from large compound libraries) would enable us to determine if FLVCR2 is responsible for the uptake of other cationic molecules into the brain; the results of such experiments could also enable the guided design of novel neurotherapeutics to promote their uptake across the BBB via a hitherto unexplored FLVCR2-mediated transport mechanism.

Our cryo-EM maps of both the OFS and IFS reveal an interesting non-protein density attached to Q214 and an additional non-protein density in the IFS map between the bound choline and residues N121, Q214, and N245 (**Extended Data Fig. 7d**). Given the proximity of these densities to key residues involved in substrate binding, and substrate itself, it is possible that they correspond to ions that either stabilise substrate binding or are co-/counter-transported with choline. Further investigations will be required to confirm their identities and elucidate the role that they and any protons may have in coordinating FLVCR2-mediated choline transport.

Given the involvement of FLVCR2 in brain angiogenesis and congenital hydrocephalus, the cardinal features of PVHH, we propose that FLVCR2-mediated choline uptake directs both primary (cell-autonomous) and secondary (non-cell-autonomous) activities. Similar to the role of FLVCR1-mediated choline uptake in other cells^12^, FLVCR2-mediated choline uptake in brain endothelial cells could support mitochondrial homeostasis and energy metabolism during cell growth, thus promoting vascular sprouting and normal brain development^32,33^. Additionally, after endothelial cells take up choline from the circulation, they must subsequently export a portion of it into the brain so that it can be utilised for phospholipid, betaine, and acetylcholine biosynthesis. Further studies are needed to investigate how choline transported by FLVCR2 is involved in metabolic and/or epigenetic signalling in endothelial cells, as well as how this molecule is subsequently transported out of endothelial cells and into the brain parenchyma. Such findings would enhance our mechanistic understanding of the cellular crosstalk in brain microenvironments during development and in disease settings, genetic or otherwise.

In closing, we have discovered that the previously-orphaned transporter FLVCR2 mediates uptake of choline across the BBB and have provided insights into the molecular mechanism by which this transport occurs. Our findings highlight the importance of the essential nutrient choline in the development of brain vasculature and the maintenance of a healthy central nervous system. Finally, as we further our understanding of FLVCR2 substrate specificity and of the system in general, this transporter could provide a means for the rational design and targeted delivery of pharmaceuticals into the brain.

## Methods

### Mice

Mice with an endogenously hemagglutinin (HA) tag on the C-terminus *of Flvcr2 (Flvcr2^HA^) w*ere generated by Biocytogen. Briefly, the targeting vector was constructed (**Extended Data Fig. 2a**), linearised, and transfected into C57BL/6J embryonic stem cells (Beijing Biocytogen Co.) by electroporation. G418-resistant embryonic stem cell clones were screened for homologous recombination by PCR. Correctly targeted clones were confirmed by Southern blot analysis and sequencing (not shown). Two positive clones were injected into BALB/c blastocysts and implanted into pseudo-pregnant females to generate chimeric mice. Chimeric mice were bred with C57BL/6J mice to obtain F1 mice carrying the *Flvcr2^HA^* allele. These mice were then bred to homozygosity. Note that homozygous breeding had no effect on fertility, frequency of live birth, or health of mice, indicating no apparent effect of attaching a HA tag to *Flvcr2* on the protein’s function. Mice with a conditional *Flvcr2^fl^* allele^34^ were a kind gift of Jianzhu Chen, Massachusetts Institute of Technology, Cambridge, MA. Animals for endothelial cell specific conditional knockout of *Flvcr2* animals were generated by crossing *Flvcr2^fl^* mice with *Cdh5Cre^ERT^*^2^ mice, a kind gift from Ralf Adams at the Max Planck Institute for Molecular Biomedicine, Muenster, Germany. We utilised mice with a conditional *Rpl22^HA^* allele (JAX Strain #011029) to identify cellular recombinants (**Extended Data Fig. 2b**). High degrees of cellular recombination were detected based on *Cre^ER^* recombination reporters and whole-brain quantitative PCR (**Extended Data Fig. 2c**). All mice were maintained on a C57BL6 background in specific pathogen-free conditions at the University of California, San Francisco (UCSF). All animal studies were followed by the protocol approved by the Institutional Animal Care and Use Committee (IACUC) at UCSF. Animals were housed in ventilated caging on a standard light/dark cycle with food and water provided ad libitum.

### Tamoxifen treatment

At 6-8 weeks of age, male and female mice were intraperitoneally injected with 75 mg per kg of body weight of tamoxifen (Sigma Aldrich) dissolved in corn oil per day for five consecutive days. Of note, mice with conditional endothelial cell specific knockout of *Flvcr2* in adulthood showed no obvious vascular abnormalities or dysfunction^16^.

### Immunohistochemistry

Adult mice (*Flvcr2^HA^*, *Flvcr2^Fl/+^Cdh5Cre^ERT2^* (control) and *Flvcr2^Fl/Fl^Cdh5Cre^ERT2^* (*Flvcr2* conditional knockout, cKO) were harvested and perfused transcardially with PBS and then with 4% paraformaldehyde (PFA). The brains were fixed overnight in 4% PFA, transferred to 30% sucrose overnight, and frozen in OCT. Sagittal sections (20 μm) were obtained in a cryostat and mounted onto glass slides. The sections were further processed for immunohistochemistry. Briefly, the sections were blocked and permeabilised for 1 hour in PBS containing 0.5% Triton X-100, 1% BSA, and 5% donkey serum (blocking buffer). Then, the sections were incubated with primary antibodies against rabbit anti-HA (Cell Signaling Technology), rat anti-ICAM2 (BD Biosciences), goat anti-CD31 (R&D Systems), diluted goat anti-collagen IV (Southern Biotech), and/or rabbit anti-collagen IV (AbD Serotec) diluted in blocking buffer in a humidified chamber overnight at 4°C. The next day, the slides were washed three times with 1X PBS. The sections were then incubated with corresponding secondary antibodies (Alexa Fluor 405, 488, 568, Cy3 or 647, Life Technologies) in blocking buffer for two hours at room temperature. The slides were washed again three times with 1X PBS and mounted with Prolong Gold Antifade Mounting media (Invitrogen). The images were taken using a motorised Zeiss 780 upright laser-scanning confocal microscope with either 20X/1 water immersion or 63X/1.4 oil immersion objective (Zeiss). Raw images were analysed using FIJI/ImageJ (NIH).

### *In vivo* choline uptake studies

Choline uptake experiments were performed seven days after tamoxifen-mediated gene deletion. [Methyl-^3^H]-Choline Chloride (Perkin Elmer) (0.3 mCi/kg) was administered intravenously through retro-orbital injection in 0.1 ml saline and allowed to circulate for 30 min. The specific activity of the [^3^H]ChCl Choline was 80.33 Ci/mmol. 30 min after injection, the mice were killed. Blood samples were collected, and serum isolated by centrifuging blood samples for 5 min at 2000 x g. The brains were collected and washed in ice cold PBS then dissolved in solubilising buffer (1M KOH) at 65°C. The radioactive [^3^H]-Choline accumulation in the brain was measured using Scintillation counter (Perkin Elmer). Total brain accumulation of radioactivity was calculated by percentage accumulation (DPM per gram of brain / DPM per ml of serum) and then normalised to control.

### Cell culture

Both BEND3.1 and HBEC5i cell lines were purchased from American Type Culture Collection (ATCC). The two BEND3.1 *Flvcr2* knock out (KO) cell lines were generated using CRISPR method (**Extended Data Fig. 2b**; Synthego). The KO1 was generated by addition of two bases, while the KO2 line was generated by deleting one base pair. Both the cell lines were validated by Sanger Sequencing. BEND3.1 cells were cultured in Dulbecco’s modified Eagle medium (DMEM; ATCC) with high glucose (4.5 g/litre), sodium bicarbonate (1.5 g/litre), 4 mM L-glutamine, 1 mM sodium pyruvate and 10% foetal bovine serum (FBS; Thermo Fisher Scientific). For HBEC5i, the cell culture plates were coated with 0.1% Gelatin (ATCC) for an hour at 37°C. The cells were cultured in DMEM:F12 (ATCC) containing 2.5 mM L-Glutamine, 15 mM HEPES, 0.5 mM Sodium pyruvate, 1200 mg/litre sodium bicarbonate and 10% FBS. Parental HEK293 cells were purchased from either the UCSF Cell and Genome Engineering core facility or ATCC. HEK293 cells were grown in RPMI 1640 medium (Gibco) supplemented with 10% FBS. HEK293 *Flvcr1* KO cells were generated by CRISPR-Cas9 gene editing as described previously^12^. *Flvcr1* KO HEK293 cells with stable expression of human *Flvcr1 (Flvcr1* OE) were generated as described previously^12^. Both HEK293 *Flvcr1* KO and *Flvcr1* KO *+ Flvcr1* OE were a kind gift from the Kivanc Birsoy laboratory, Rockefeller University, New York, NY, USA. These cells were cultured in RPMI 1640 medium supplemented with 10% FBS. All cells were maintained in an incubator maintained at 37°C, 5% CO2, and 80% relative humidity (RH).

### Knock-down of *Flvcr2* in HBEC5i cells

Human *Flvcr2* was knocked down in HBEC5i cells using siRNA (Silencer Select siRNA; Ambion). Cells were transfected with either scrambled siRNA or *Flvcr2* siRNA using Lipofectamine 3000 (Fisher Scientific). *Flvcr2* knock-down was confirmed by qPCR 48 hours after transfection (**Extended Data Fig. 2d**).

### Transfection of *Mm*FLVCR2 cDNA in HEK293 cells

Both parental and *Flvcr1* KO HEK293 cells were transfected with GFP-tagged *Flvcr2* (*Mm*FLVCR2 cDNA; *Flvcr2* OE) using Lipofectamine 3000. Approximately 10,000-20,000 cells were seeded in a 48 well plate. Briefly, 2 µg DNA diluted in 250 µL of OptiMEM (Fisher Scientific) containing p3000. The diluted DNA was then added to diluted Lipofectamine 3000 in 250 µL of OptiMEM. The DNA-Lipofectamine complex was incubated for 20 min at room temperature. The mixture was then added to the cells dropwise. All experiments were performed 48 hours post-transfection.

### Cell-based radiolabelled choline uptake experiments

Cells were seeded at a density of 30,000 cells per 48 well plate in triplicate. On the day of experiment, the cells were incubated in Krebs-Ringer Buffer (120 mM NaCl, 5 mM KCl, 2 mM CaCl_2_, 1 mM MgCl_2_, 25 mM NaHCO_3_, 5.5 mM HEPES, 1 mM D-Glucose; pH 7.2) for 30 min. Cells were then incubated with 20 nM choline chloride in Krebs-Ringer Buffer for 30 min at 37°C. The specific activity of the choline was 80.33 Ci/mmol. For all concentrations of ChCl, 0.093% of the total choline was [^3^H]ChCl. The cells were then washed twice with ice cold Krebs-Ringer Buffer. Next, the cells were solubilised with 750 uL of solubilising buffer (1% SDS; 0.2 N NaOH) for an hour at room temperature on an orbital shaker. 690 uL of the samples were transferred to a scintillation vial with 2 mL of Insta-Gel Plus scintillation cocktail (Perkin Elmer). Radioactivity was measured with the Scintillation counter (Perkin Elmer). The remaining samples were used to calculate protein concentration. Radioactivity was normalised to the protein concentration of each well. The data were represented as relative fold change to control. For BEND3.1 cells, parental cells (wild type; WT) control and *Flvcr2* knockout (KO1 and KO2) cells were seeded and incubated overnight and then the uptake experiment performed the following day. For HBEC5i cells, *Flvcr2* was knocked down using siRNA. Uptake experiments were performed 48 hours after siRNA transfection. For HEK293 cells, mouse *Flvcr2* was overexpressed by transient transfection with *Mm*FLVCR2 cDNA (*Flvcr2 OE*). The uptake experiments were performed 48 hours post transfection.

### RNA isolation for *in vitro* studies

Total RNA from cells was isolated using TRI reagent following manufacturer’s instructions. HBEC5i cells with treated with either non-specific siRNA or *Flvcr2* siRNA were harvested and kept in 1 mL TRI reagent (Qiagen).

### RNA isolation for *in vivo* studies

Brains were collected from either *Flvcr2^Fl/+^Cdh5Cre^ERT2^*(control) and *Flvcr2^Fl/Fl^ Cdh5Cre^ERT2^* (*Flvcr2* cKO) adult mice and kept in TRI reagent (Qiagen) and RNA was isolated following manufacturer’s instructions. Briefly, about 100 mg of tissue samples were homogenised in 1 mL of TRI reagent at room temperature using a tissue lyser (Qiagen). The homogenised samples were transferred to fresh microcentrifuge tubes for RNA isolation. Both *in vivo* and *in vitro* samples were incubated in TRI reagent for 10 min at room temperature. 200 μL chloroform was then added, mixed, and incubated for 10 min at room temperature. The samples were next centrifuged at 12,000 x g for 15 min at 4°C. The upper aqueous phase containing the RNA was then collected in a separate tube and mixed with 500 μL of isopropyl alcohol incubated for 10 min. RNA was precipitated by centrifugation at 12,000 x g for 10 min at 4°C. RNA pellet obtained was washed with 75% ethanol by and dissolved in 40 μL of RNase free DEPC water. RNA was quantified by spectrophotometric reading at 260nm, and purity was checked at 260/280 nm using a Nano Drop (Fisher Scientific).

### RNA sequencing

Total RNA (RIN > 6) was extracted from 1 x10^6^ BEND3.1 or HBEC5i cells using the QIAGEN RNeasy Micro kit and used for PolyA+ unstranded library synthesis with NEBNext Ultra II RNA Library Prep kit. Sequencing was performed with an Illumina HiSeq4000 instrument in a 150 pb paired-end setup. Demultiplexed FASTQ files were aligned to the mouse (mm10) or human (hg38) genome with Rsubread, quantified with FeatureCounts, and normalised with DESeq2 procedure. Heatmaps were constructed with the pheatmap package. Raw data from BEND3.1 and HBEC5i cells were deposited in GEO under accession GSE244349. Other datasets are as follows: iPSC neurons and HEK293 are from GSE219195; HeLa cells are from GSE228126; specific cell types from human brain are from GSE73721; specific cell types from mouse brain are from GSE52564.

### Small-scale expression screen of FLVCR2 variants

Initial expression studies were performed with six FLVCR2 variants: *Homo sapiens* (*Hs*FLVCR2; NCBI – NP_060261.2), *Mus musculus* (*Mm*FLVCR2; NCBI – NP_663422.1) *Eptesicus fuscus* (*Ef*FLVCR2; NCBI – XP_027997147.2), *Canis lupus dingo* (*Cld*FLVCR2; NCBI – XP_025299773.1), *Bos taurus* (*Bt*FLVCR2; NCBI – NP_001179072.1), and *Danio rerio* (*Dr*FLVCR2; NCBI – XP_693589.3). To express these in mammalian cells, the sequences were cloned into pFM1.2^35^ as GFP fusions with a decahistidine affinity tag and flag affinity (DYKDDDDK) tag at the 3′ end of the gene. In brief, 5 μg of each construct was diluted into 250 μL OptiMEM (Thermo Fisher Scientific) and mixed with 250 μL OptiMEM containing 20 μg of polyethylenimine (PEI – maximum molecular mass of 40,000 Da; Polysciences). This mixture was added to 20.0 × 10^6^ Expi293F GnTI-Cells (Thermo Fisher Scientific) in a total volume of 10 ml. The cell transfection mixtures were then incubated at 37°C for 72 h with 5% CO_2_, 70% humidity, and shaking set at 115 rpm. Transfected cells were then collected, centrifuged at 800 x *g* for 10 min at 4°C and washed once on ice in 1X PBS. Each pellet was resuspended and solubilized in 20 mM HEPES pH 7.5, 200 mM NaCl, 20 mM MgSO_4_, 0.5 mM phenylmethylsulfonyl fluoride (PMSF), cOmplete EDTA-free protease inhibitor cocktail (Roche), 10 μg ml^−1^ DNase I (Roche) and 8 μg ml^−1^ RNase (Sigma-Aldrich) supplemented with 1% (w/v) *n*-dodecyl-β-d-maltopyranoside (DDM) and 0.1% (w/v) cholesteryl hemisuccinate (CHS) at 4°C for 2 h. Insoluble material was removed by ultracentrifugation in a single-angle rotor at 4°C for 45 min. To evaluate expression and stability of the GFP-tagged protein, the solubilised supernatants were then subjected to fluorescence-coupled size-exclusion chromatography (FSEC)^36^ using a Superdex 200 Increase 5/150 GL column (Cytiva) attached to an Agilent Technologies 1200 Series HPLC system with an in-line fluorescence detector. These studies identified FLVCR2 from *Mus musculus* (*Mm*FLVCR2) and *Bos taurus* (*Bt*FLVCR2) as having the best yield and mono-dispersity (**Extended Data Fig. 3a**). Given that *Mus musculus* is an established system for studying FLVCR2^37,38^, and that *Mm*FLVCR2 shares 87% identity with *Hs*FLVCR2, we chose to move forward with *Mm*FLVCR2 for our structural and functional experiments.

### FLVCR2 protein expression and purification

The *Mm*FLVCR2 full-length open-reading frame was cloned into pBacMam (Thermo Fisher Scientific) using ligation-independent cloning^39^. *Mm*FLVCR2 was fused at its 3′ end with a tobacco etch virus protease cleave site (ENLYFQ/S) and a decahistidine affinity tag followed by a flag affinity tag. The resulting plasmid was transformed into *E. coli* DH10Bac competent cells to generate the baculovirus DNA using a bac-to-bac protocol (Thermo Fisher Scientific). Recombinant P1 baculovirus DNA was transfected into and cultured in Sf9 cells (Expression System) in penicillin-streptomycin-free ESF 921 protein-free insect cell culture media (Expression Systems) in the presence of PEI. Virus passages were then amplified in penicillin-streptomycin-containing media (Roche) until cells became visibly swollen and demonstrated reduced viability statistics. For protein expression, 8 ml P4 virus was used to infect 800 ml cultures of Expi293F GnTI-Cells (Thermo Fisher Scientific) cells at 2-3 × 10^6^ cells supplemented with 1% Fetal Bovine Serum (FBS). Following transduction, the cells were then grown in a shaking incubator (115 rpm) at 37°C for 72 h with 5% CO_2_ and 70% humidity before harvesting as described above.

Cell pellets were homogenised in low-salt buffer (10 mM HEPES pH 7.5, 10 mM KCl, 10 mM MgCl_2_, 0.5 mM PMSF, cOmplete EDTA-free protease inhibitor cocktail, 10 μg mL^−1^ DNase I, and 8 μg mL^−1^ RNase) in a glass homogeniser. Membrane fractions were isolated by ultracentrifugation at 40,000 x *g* in a type 45 Ti rotor (Beckman Coulter). Membrane fractions were further homogenised and washed twice with high-salt buffer (10 mM HEPES pH 7.5, 10 mM KCl, 10 mM MgCl_2_, 1 M NaCl, 0.5 mM PMSF, cOmplete protease inhibitor cocktail, 10 μg mL^−1^DNase I and 8 μg mL^−1^ RNase) in a glass homogeniser followed by ultracentrifugation. The washed membrane fractions were resuspended again by homogenising in buffer comprising 20 mM HEPES pH 7.5, 200 mM NaCl and 0.5 mM PMSF and cOmplete protease inhibitor cocktail and stored at −80°C until use.

To solubilise the membrane fraction, the pellet was resuspended in 4x the cell pellet weight of solubilisation buffer (20 mM HEPES pH 7.5, 200 mM NaCl, 0.5 mM PMSF, cOmplete protease inhibitor cocktail, 10 μg mL^−1^ DNase I and 8 μg mL^−1^ RNase) containing DDM with CHS in a 10:1 (w/w) ratio to a final concentration of 1% (w/v) detergent and incubated at 4°C for 2 h with gentle agitation. Insoluble material was removed by ultracentrifugation at 40,000 x *g* in a type 45 Ti rotor at 4°C for 30 min. The supernatant was then incubated with pre-equilibrated Ni^2+^-NTA resin (Qiagen) in the presence of 30 mM imidazole and left at 4 °C overnight with gentle rotation. The resin was washed with 10 column volumes of buffer comprising 20 mM HEPES pH 7.5, 200 mM NaCl, 60 mM imidazole, 0.1% DDM, and 0.01% CHS. Bound protein was eluted with 3-5 column volumes of buffer consisting of 20 mM HEPES pH 7.5, 200 mM NaCl, 200 mM imidazole, 0.05% DDM, and 0.005% CHS. For experiments where detergent-purified *Mm*FLVCR2 was the desired end product, the eluted product was then further purified by loading on a Superdex 200 Increase 10/300 GL size-exclusion chromatography (SEC) column (Cytiva) in gel-filtration buffer (20 mM HEPES pH 7.0, 150 mM NaCl, 0.025% DDM, and 0.0025% CHS). SEC fractions containing monomeric *Mm*FLVCR2 were then pooled and concentrated using 100 kDa MWCO Amicon centrifugal filtration units (Millipore Sigma) as required for subsequent experiments.

### Amino acid sequence alignments

To analyse the sequence conservation across FLVCR2 and FLVCR1 variants, we aligned four FLVCR2 and four FLVCR1 amino acid sequences using Clustal Omega^40^. The sequences corresponded to *Homo sapiens* (*Hs*FLVCR2; NCBI – NP_060261.2), *Mus musculus* (*Mm*FLVCR2; NCBI – NP_663422.1), *Canis lupus dingo* (*Cld*FLVCR2; NCBI – XP_025299773.1), *Bos taurus* (*Bt*FLVCR2; NCBI – NP_001179072.1), *Homo sapiens* (*Hs*FLVCR1; NCBI – NP_054772.1), *Mus musculus* (*Mm*FLVCR1; NCBI – NP_001074728.1), *Canis lupus familiaris* (*Clf*FLVCR1; NCBI – XP_003639227.1), and *Bos taurus* (*Bt*FLVCR1; NCBI – NP_001192948.1). A second multiple sequence alignment was also performed to map sequence conservation across mammalian FLVCR2 orthologs on the structure of *Mm*FLVCR2 (**Extended Data Fig. 4**). To select sequences for alignment, the *Mm*FLVCR2 protein sequence was used to query mammalian RefSeq proteins, from which the top 100 hits were used.

### Proteoliposomes preparation

Detergent purified *Mm*FLVCR2 was reconstituted into preformed liposomes composed of brain polar lipids (Avanti Polar Lipids, Inc.) at a protein-to-lipid ratio of 1:150 (w/w) following a previously-described protocol^41^. The lumen of the proteoliposomes was composed of 100 mM Tris/Mes, pH 7.5 or 100 mM KP_i_ pH 7.5. To maintain their stability, the proteoliposomes were aliquoted, rapidly frozen in liquid nitrogen, and stored at −80°C. In all experimental setups, liposomes with the same composition but lacking the FLVCR2 protein were used as a control.

### Liposome-based radiolabelled choline uptake assays

Uptake of 0.1 and 1 µM [^3^H]ChCl (5 Ci/mmol, American Radiolabeled Chemicals, Inc.) was performed in proteoliposomes (∼ 50 ng *Mm*FLVCR2 per transport assay) in 50 μL assay buffer composed of 100 mM Tris/Mes, pH 5.5-8.5 in the presence or absence of the indicated compounds. The reactions were quenched by the addition of ice-cold 100 mM KP_i_, pH 6.6, 100 mM LiCl and filtered through 0.45 µm nitrocellulose filters (Millipore). Dried filters were incubated in scintillation cocktail, and the retained radioactivity on the filters was determined by scintillation counting (Hidex, SL300). The specific uptake activity of *Mm*FLVCR2 was determined by subtracting the background d.p.m. signal determined in control liposomes (lacking *Mm*FLVCR2) from the d.p.m. measured in *Mm*FLVCR2-containing proteoliposomes. The nonspecific interaction of all compounds used in the assays with the nitrocellulose filters was determined by performing filtration experiments in the absence of (proteo)liposomes. These values (determined for each experiment) were negligible (below the signals measured with control liposomes) and were not used in the calculations. Nonlinear regression analysis of the data was performed by fitting the data to the Michaelis–Menten equation in Graph-Pad Prism 10.

### Binding studies

Experiments used 1 µM [^3^H]ChCl (80 Ci/mmol). Scintillation proximity-based assay (SPA) binding of 25 ng of *Mm*FLVCR2 was assayed in buffer composed of 20 mM Tris/Mes pH 8.5, 200 mM NaCl, 0.025% DDM, and 0.0025% CHS, in the presence of 1.5 mg/mL of His tag SPA beads, and in the presence or absence of indicated reagent. Then, 800 mM imidazole, which competes with the His tag (fused to *Mm*FLVCR2) for binding to the copper-coated SPA beads, was used to determine the non-proximity background signal.

### FLVCR2 nanodisc reconstitution

Following elution from the NiNTA resin (as described above), imidazole was removed from the eluted protein using a PD10 desalting column (Cytiva). The *Mm*FLVCR2 protein was them reconstituted into nanodiscs using a 1:200:5 molar ratio of protein: 1-palmitoyl-2-oleoyl-*sn*-glycero-3-phospho-(1′-rac-glycerol) (POPG): membrane-scaffold protein 1D1 (MSP1D1). This mixture was then incubated at 4°C for 2 h with gentle agitation before reconstitution was initiated by removing detergent by incubating with Bio-beads (Bio-Rad) at 4°C overnight with constant rotation. Bio-beads were then removed and the nanodisc reconstitution mixture was bound again to Ni^2+^-NTA resin at 4°C for 2 h in the absence of imidazole to remove empty nanodiscs. The resin was washed with 6 column volumes of wash buffer (20 mM HEPES pH 7.5, 200 mM NaCl and 50 mM imidazole) followed by 4 column volumes of elution buffer (20 mM HEPES pH 7.5, 200 mM NaCl and 300 mM imidazole). The eluted protein was further purified by loading on a Superdex 200 Increase 10/300 GL SEC column (Cytiva) in gel-filtration buffer (20 mM HEPES pH 7.0, 150 mM NaCl). SEC fractions containing monomeric nanodisc-reconstituted *Mm*FLVCR2 were then pooled and concentrated using 100 kDa MWCO Amicon centrifugal filtration units (Millipore Sigma) as required for subsequent experiments.

### Identification of FLVCR2-specific Fabs and initial binding validation

*Mm*FLVCR2 was reconstituted into nanodiscs using chemically biotinylated MSP1D1 as described above and used for biopanning against phage-display Fab Library E^42^. The efficiency of biotinylation was subsequently evaluated by capture using streptavidin-coated paramagnetic particles (Promega). The first round of selection was performed using 400 nM of nanodisc-reconstituted *Mm*FLVCR2 diluted in selection buffer (20 mM HEPES pH 7.5, 150 mM NaCl, 1% BSA) using a previously described protocol^25,43,44^. Four additional rounds of selection were performed as described with decreasing target concentration (200 nM, 100 nM, 50 nM, and 20 nM) using the magnetic particle purification system KingFisher (Thermo Fisher Scientific). In every subsequent round, the amplified phage pool from the previous round was used as the input. Prior to being used for selection, each phage pool was precleared by incubating with 100 µL of streptavidin paramagnetic beads. Additionally, in all rounds, 5-10 times molar excess of nonbiotinylated MSP1D1 nanodiscs were added to the selection buffer as competitors to reduce the presence of non-specific binders. In rounds two to five, selection buffer supplemented with 1% Fos-choline-12 was used to release the target and bound phages from the nanodiscs. Cells infected after the last round were plated on LB agar with ampicillin (100 µg/mL) and phagemids from individual clones were subjected to sequencing. Single-point phage ELISA was used to validate the specificity of identified unique binders as described previously^43,44^.

*Mm*FLVCR2-specific binders were sequenced at the University of Chicago Comprehensive Cancer Center Sequencing Facility and unique clones were subsequently sub-cloned into Fab expression vector RH2.2 (kind gift of S. Sidhu)^43^. Sub-cloned Fabs were sequence-verified, expressed, and purified as previously described^43^. Purified Fabs were additionally validated using affinity estimation ELISA to quantitatively assess binding to *Mm*FLVCR2^25^.

### Epitope Masking ELISA to identify additional epitopes

Epitope masking single-point phage ELISA was performed using a protocol adapted from previous work^45^ for the 40 Fab-phage clones that produced the highest ELISA signal in the earlier experiment. Binding to *Mm*FLVCR2 was tested in the presence and absence of 1 uM purified Fab 32, which was used as an epitope masking reagent. Fab 32 was incubated with *Mm*FLVCR2 prior to addition of the phage solution, and was added to one set of the phage dilutions for competition. The remainder of the ELISA protocol was performed as described above and the signal was detected using the same protocol. Wells containing only BSA were used as negative controls.

### Mass photometry (MP, iSCAMS)

Mass photometry (MP) tests were conducted utilizing a Refeyn OneMP (Refeyn Ltd). For data acquisition, we used AcquireMP (version 2023 R1; Refeyn Ltd). The specimens were examined on microscope slides with glass thickness of 1 ½ 24 x 50 mm (Corning), which were cleaned using deionised water and isopropanol. Prior to carrying out measurements, silicone moulds were positioned atop the slide to create reaction chambers. Calibration of the device was achieved using NativeMark Protein Standards (NativeMark™ Unstained Protein Standard, Thermo Fisher). 10 μL of room-temperature buffer was dispensed into a designated well and the focal location was identified. 1 μL of the protein with an end concentration post-dilution of 200 nM was incorporated into the well and mixed thoroughly. MP readings spanned 60 seconds, ensuring the observation of a minimum of 2 x 10^3^ discrete binding occurrences. Analysis was conducted through the DiscoverMP software.

### Preparation of FLVCR2-Fab complexes for structural analysis

Detergent-purified and nanodisc-reconstituted *Mm*FLVCR2 were incubated with Fabs on ice for 2 h using a 1:1.5 molar ratio of protein to Fab. The *Mm*FLVCR2–Fab complexes were then concentrated to 500 µL, filtered, and separated by SEC using a Superdex 200 Increase 10/300 GL column (Cytiva) as described above with gel filtration buffers supplemented with 1 mM ChCl. While Fabs were originally identified using nanodisc-reconstituted *Mm*FLVCR2, our initial single-particle analyses of Fab-complexed, nanodisc-reconstituted *Mm*FLVCR2 demonstrated that these particles adopted a highly preferred orientation (lacking side-views) which could not be overcome by the addition of amphipathic molecules such as amphipol, CHAPS detergent, fluorinated octyl maltosides, or glycyrrhizic acid. On the other hand, side views were abundant in detergent-purified *Mm*FLVCR2-Fab particles, thus for our high-resolution structures we chose to use Fab-complexed, detergent-purified *Mm*FLVCR2.

### Single-particle cryo-EM vitrification and data acquisition

For the outward-facing state (OFS) structure, detergent-purified *Mm*FLVCR2 was complexed with Fab FLV23 as described above, whereas Fab FLV9 was the Fab of choice for the inward-facing state (IFS) structure, given that this Fab bound at a distinct epitope. Detergent-purified *Mm*FLVCR2-Fab complexes were purified by SEC and concentrated to 5.5-8 mg ml^−1^ using a 100 kDa MWCO concentrator (Amicon). The concentrated sample was then frozen using a Vitrobot (Thermo Fisher Scientific) by applying 3 μL of it to plasma-cleaned 0.6/1 μm holey gold UltrAuFoil grids (Quantifoil). Following a 30 s incubation, the grids were blotted for 4.5-5 s (blot force of 3) using 595 filter paper (Ted Pella, Inc) before being immediately plunged into liquid ethane for vitrification. The plunger was operated at 4°C with greater than 90% humidity to minimise evaporation and sample degradation.

For both structures, movies were recorded using a Titan Krios Cryo-Transmission Electron Microscope (Thermo Fisher Scientific) at the Columbia University Cryo-Electron Microscopy Center, operating at 300 kV, and equipped with an energy filter and a K3 direct electron detection filter camera (Gatan K3-BioQuantum) using a pixel size of 0.83 Å. An energy filter slit width of 10 eV was used during the collection and was aligned automatically every hour using the Leginon software package^46^. Data collection was performed using a dose of 58.2 e^−^ Å^−2^ across 50 frames (50 ms per frame) at a dose rate of approximately 16.0 *e*^−^ per pixel per second, using a set defocus range of −0.8 μm to −1.5 μm. A 100 μm objective aperture was used. In total, 10,073 movies were collected for the OFS structure and 16,402 movies were collected for the IFS structure.

### Single-particle cryo-EM data processing

Unless otherwise stated, all cryo-EM data processing was performed using CryoSPARC v.4.1.2^47^. Movie frames were aligned using Patch Motion Correction (B-factor of 500) after which Contrast Transfer Function (CTF) estimations were made using the Patch CTF estimation tool. Micrographs were then filtered based on CTF estimations, defocus, total frame motion, and ice thickness which returned 6,247 and 11,703 micrographs for the *Mm*FLVCR2-Fab FLV23 complex (OFS) and *Mm*FLVCR2-Fab FLV9 complex (IFS), respectively.

For the *Mm*FLVCR2-Fab FLV23 complex (OFS), an initial round of particle picking was performed using the blob picker tool (diameter 100-280 Å), followed by iterative 2D and 3D classifications to generate 2D class average templates and 3D volumes for *Mm*FLVCR2+Fab, *Mm*FLVCR2 alone, and high-noise decoy classes. Particle picking was then repeated using TOPAZ^48^ as implemented in CryoSPARC. A total of 1,976,209 particles were extracted with a box size of 360 pixels and 4x binning. These particles were sorted using one round of 2D-classification to remove high-contrast noise particles, after which the selected 1,927,624 particles were fed into two iterative rounds of heterogeneous refinement using 8-13 volumes of *Mm*FLVCR2+Fab, *Mm*FLVCR2 alone, and high-noise decoy classes (previously generated from the initial Blob Picking) as input volumes. The highest resolution class from the second heterogeneous refinement contained 466,906 particles, which were then further sorted by one round of 2D-classification to yield a total of 433,845 particles which were then re-extracted with a box size of 360 pixels (no binning). Next, global CTF correction per image group was performed^47,49^, and the particles were then used as input for a non-uniform refinement. Local refinement was then performed with a mask around the *Mm*FLVCR2 portion of the particle, using an initial lowpass resolution of 8 Å, searching over a range of 9 degrees in orientations and 3 Å in shifts to yield a 2.49 Å resolution map (**Extended Data Fig. 5**).

For the *Mm*FLVCR2-Fab FLV9 complex (IFS), 4,886,381 particles were picked using the blob picker tool (diameter 100-250 Å) and extracted with a box size of 320 pixels and 4x binning. Following iterative 2D-classification, 745,711 particles were re-extracted with a box size of 320 pixels (no binning) and fed into a three-class *ab initio* reconstruction. The highest resolution volume from this class was then used as input for a heterogeneous refinement with six classes using 3D volumes of *Mm*FLVCR2+Fab, *Mm*FLVCR2 alone, and high-noise decoy classes generated through 2D classification and *ab initio* reconstruction. The highest resolution class from this refinement contained 308,791 particles which were then fed into a non-uniform refinement. Global CTF correction per image group was then performed^47^, and particles were re-extracted using a box size of 380 pixels (no binning). Local refinement was then performed with a mask around the *Mm*FLVCR2 portion of the particle, using an initial lowpass resolution of 8 Å, searching over a range of 9 degrees in orientations and 3 Å in shifts to yield a 2.77 Å resolution map (**Extended Data Fig. 9**).

### Structural model building, refinement, and analysis

To build the *Mm*FLVCR2 model, we used an initial model generated using AlphaFold^50^. The model was fit to the map in UCSF Chimera^51^ and transferred to Coot^52,53^ for manual model building. For Fab model building, the Fab portion of MFSD2A-Fab complex (PDB: 7MJS)^43^ was used as a starting template. In both the IFS and OFS structures we observed non-protein density within the central cavity that we assigned as choline. Model refinement and adjustment was performed in Coot^52,53^ until no further improvements were observed. Models were validated using the cryo-EM validation tool in Phenix^54^.

## Acknowledgements

We gratefully acknowledge the assistance of members of the Mancia and Arnold labs and the Columbia Cryo-EM facility. We are also grateful to S.W.J. Mooney for his assistance during manuscript preparation. This work was supported by NIH grants (R21 NS129105-01 to T.A., F.M. and R.J.C.). R.J.C. is supported by an NIH K99 Fellowship (HL166866-01). F.M.’s contribution is supported by NIGMS grant R35GM132120. Some of the work was performed at the Center for Membrane Protein Production and Analysis (COMPPÅ; NIH P41 GM116799 to Wayne A. Hendrickson), located at the New York Structural Biology Center.

## Author contributions

The project was conceived by R.J.C., T.A., and F.M.. Animal studies and cell-based assays were designed and performed by D.M., N.S.A., K.K., S.W.Y., and T.A.. N.S.A. and T.A. analysed expression data. SPA and liposomal uptake assays were designed and performed by E.G.I., R.J.C, T.C., and M.Q.. Cloning was performed by T.C. and B.K.. R.J.C. designed and performed expression screening experiments, produced baculovirus, and optimized protein expression and purification with assistance from T.C., A.R., and B.C.C.. S.K.E., T.G., and A.A.K. identified and purified the Fabs, and performed epitope binning assays. R.J.C. prepared the sample for structure analysis, screened and optimized sample vitrification, and generated cryo-EM data with assistance from Z.Z. and A.R.. Initial cryo-EM data sets were analysed by R.J.C. and Z.Z. High resolution cryo-EM data analysis and model building was performed by R.J.C. with guidance from O.B.C.. R.J.C., D.M., T.A., and F.M. wrote the manuscript with input from B.K.. and A.A.R.. R.J.C., D.M., T.A., B.K., and N.S.A. prepared the figures.

## Correspondence and requests for materials

Should be addressed to R.J.C., T.A., or F.M..

## Data availability

All raw movie frames, micrographs, the particle stack and relevant metadata files will be deposited into EMPIAR. The electron density map will be deposited into EMDB. The model will be deposited into PDB.

---

**Extended Data Table 1.** FLVCR2 outward-facing conformation cryo-EM data.

| *Mm*FLVCR2-FLV23 complex<br>(EMD-XXXX)<br>(PDB ID XXXX) |  |
| --- | --- |
| **Data collection and processing** |  |
| Magnification | 105,000 |
| Voltage (kV) | 300 |
| Electron exposure (e-/Å<sup>2</sup>) | 58.2 |
| Dose rate (e-/pixel/s) | 16 |
| Defocus range (µm) | 0.8 -1 .5 |
| Pixel size (Å) | 0.83 |
| Symmetry imposed | C1 |
| Initial micrographs collected (no.) | 10,073 |
| Final micrographs processed (no.) | 6,274 |
| Initial particle images (no.) | 1,927,624 |
| Final particle images (no.) | 433,845 |
| Map resolution (Å) | 2.49 |
| FSC threshold | 0.143 |
| Map sharpening *B* factor (Å<sup>2</sup>) | -58.6 |
| Residue range | 99-312 and 319-520 (FLVCR2)<br>31-100 (FLV23 vFab light chain)<br>1-129 (FLV23 vFab heavy chain) |
| Model composition |  |
| Non-hydrogen atoms | 4797 |
| Protein residues | 615 |
| Ligands | 1 |
| *B* factors (Å<sup>2</sup>) |  |
| Protein | 37.30 - 124.35 (mean = 55.79) |
| Ligand | 20.00 |
| R.m.s. deviations |  |
| Bond lengths (Å) | 0.013 |
| Bond angles (°) | 1.888 |
| Validation |  |
| MolProbity score | 1.98 |
| Clashscore | 5.21 |
| Rotamer outliers (%) | 3.67 |
| EM-Ringer Score | 3.92 |
| Ramachandran plot |  |
| Favored (%) | 96.05 |
| Allowed (%) | 3.95 |
| Disallowed (%) | 0.00 |

**Extended Data Table 2.**
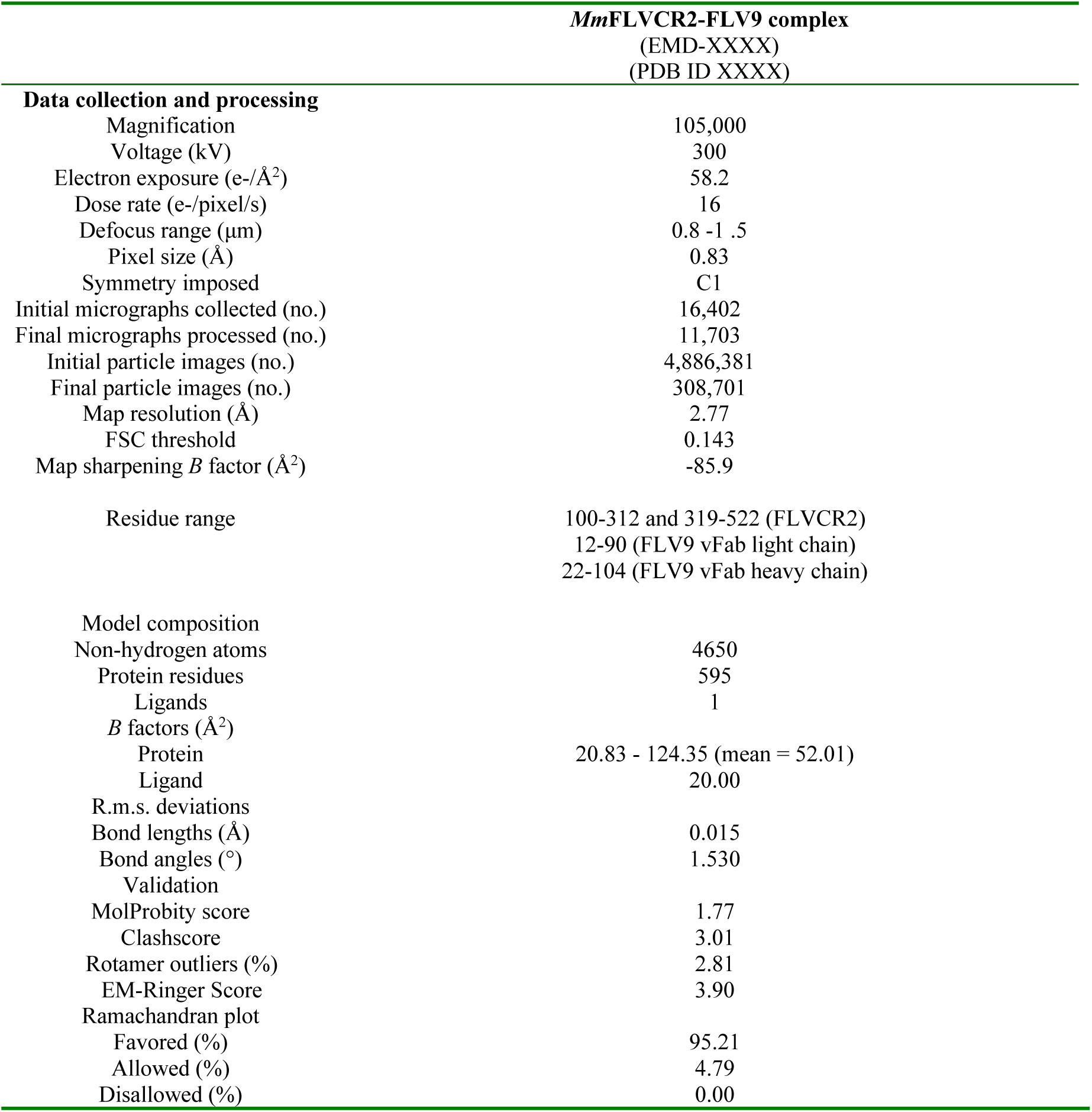
FLVCR2 inward-facing conformation cryo-EM data.

| *MmFLVCR2-FLV9 complex*<br>(EMD-XXXX)<br>(PDB ID XXXX) |  |
| --- | --- |
| **Data collection and processing** |  |
| Magnification | 105,000 |
| Voltage (kV) | 300 |
| Electron exposure (e-/Å<sup>2</sup>) | 58.2 |
| Dose rate (e-/pixel/s) | 16 |
| Defocus range (µm) | 0.8 -1 .5 |
| Pixel size (Å) | 0.83 |
| Symmetry imposed | C1 |
| Initial micrographs collected (no.) | 16,402 |
| Final micrographs processed (no.) | 11,703 |
| Initial particle images (no.) | 4,886,381 |
| Final particle images (no.) | 308,701 |
| Map resolution (Å) | 2.77 |
| FSC threshold | 0.143 |
| Map sharpening *B* factor (Å<sup>2</sup>) | -85.9 |
| Residue range | 100-312 and 319-522 (FLVCR2)<br>12-90 (FLV9 vFab light chain)<br>22-104 (FLV9 vFab heavy chain) |
| **Model composition** |  |
| Non-hydrogen atoms | 4650 |
| Protein residues | 595 |
| Ligands | 1 |
| *B* factors (Å<sup>2</sup>) |  |
| Protein | 20.83 - 124.35 (mean = 52.01) |
| Ligand | 20.00 |
| R.m.s. deviations |  |
| Bond lengths (Å) | 0.015 |
| Bond angles (°) | 1.530 |
| **Validation** |  |
| MolProbity score | 1.77 |
| Clashscore | 3.01 |
| Rotamer outliers (%) | 2.81 |
| EM-Ringer Score | 3.90 |
| **Ramachandran plot** |  |
| Favored (%) | 95.21 |
| Allowed (%) | 4.79 |
| Disallowed (%) | 0.00 |

**Extended Data Figure 1.**
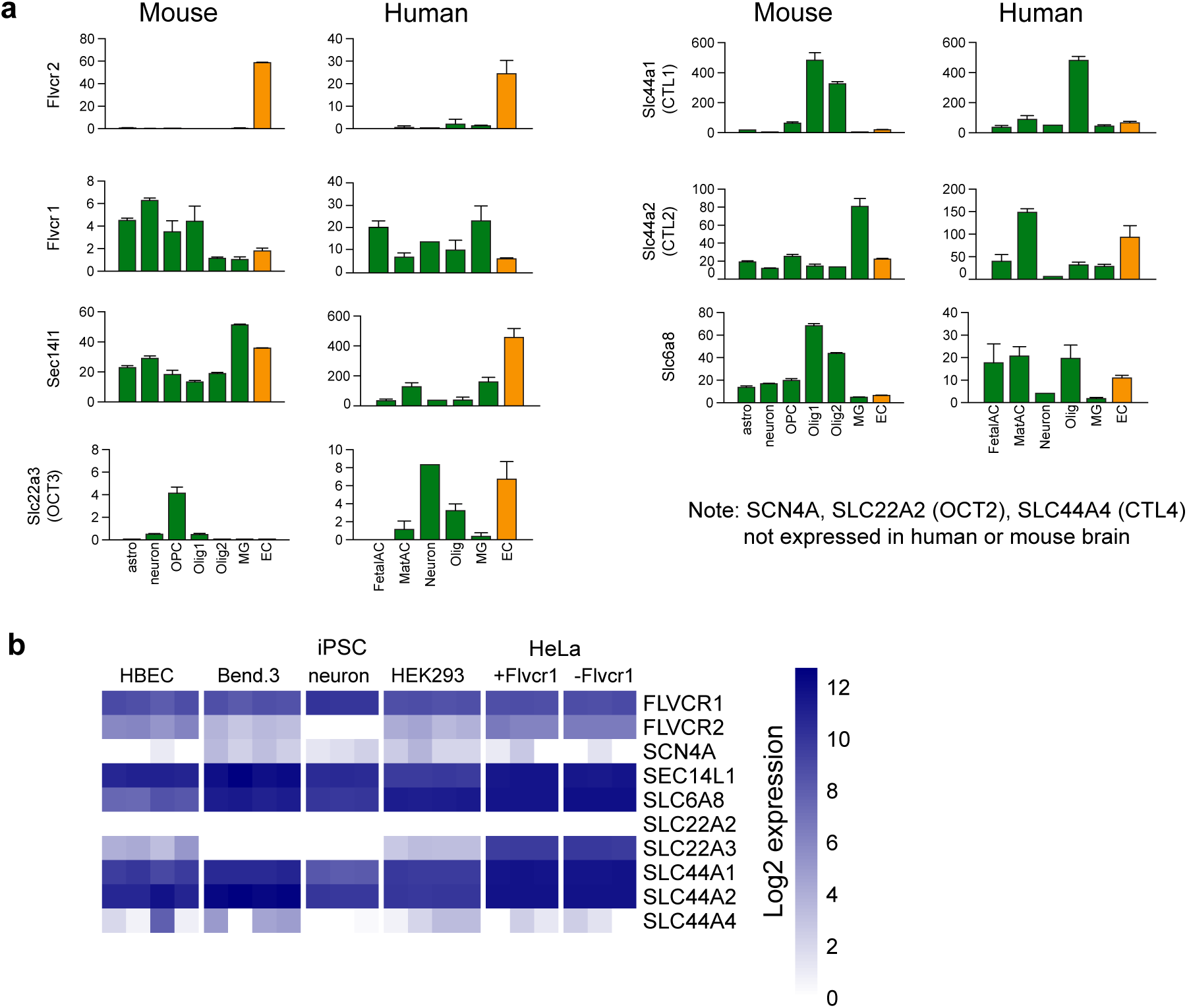
Expression profiles of *bona fide* and putative choline transporters. **a,** Expression profiles^10,11^ showing that *Flvcr2* is enriched in brain endothelial cells in both human and mouse. Other putative and validated choline transporters are not highly or specifically enriched at the BBB. Abbreviations on the horizontal axes indicate: astrocytes (astro), neuron, oligodendrocyte progenitor cells (OPC), immature oligodendrocytes (Olig1), mature oligodendrocytes (Olig2), microglia (MG), endothelial cells (EC), fetal astrocytes (FetalAC), maternal astrocytes (MatAC), and oligodendrocytes (Olig). Expression in endothelial cells is shown in orange; all other cell types are shown in green. Note the differences in vertical axis scale between transporters. **b,** RNA sequencing expression profiles from HBEC, BEND3, HEK293, human iPSC-derived neurons, and HeLa cells (with and without *Flvcr1* knockout), showing expression of putative and validated choline transporters in these cells.

**Extended Data Figure 2.**
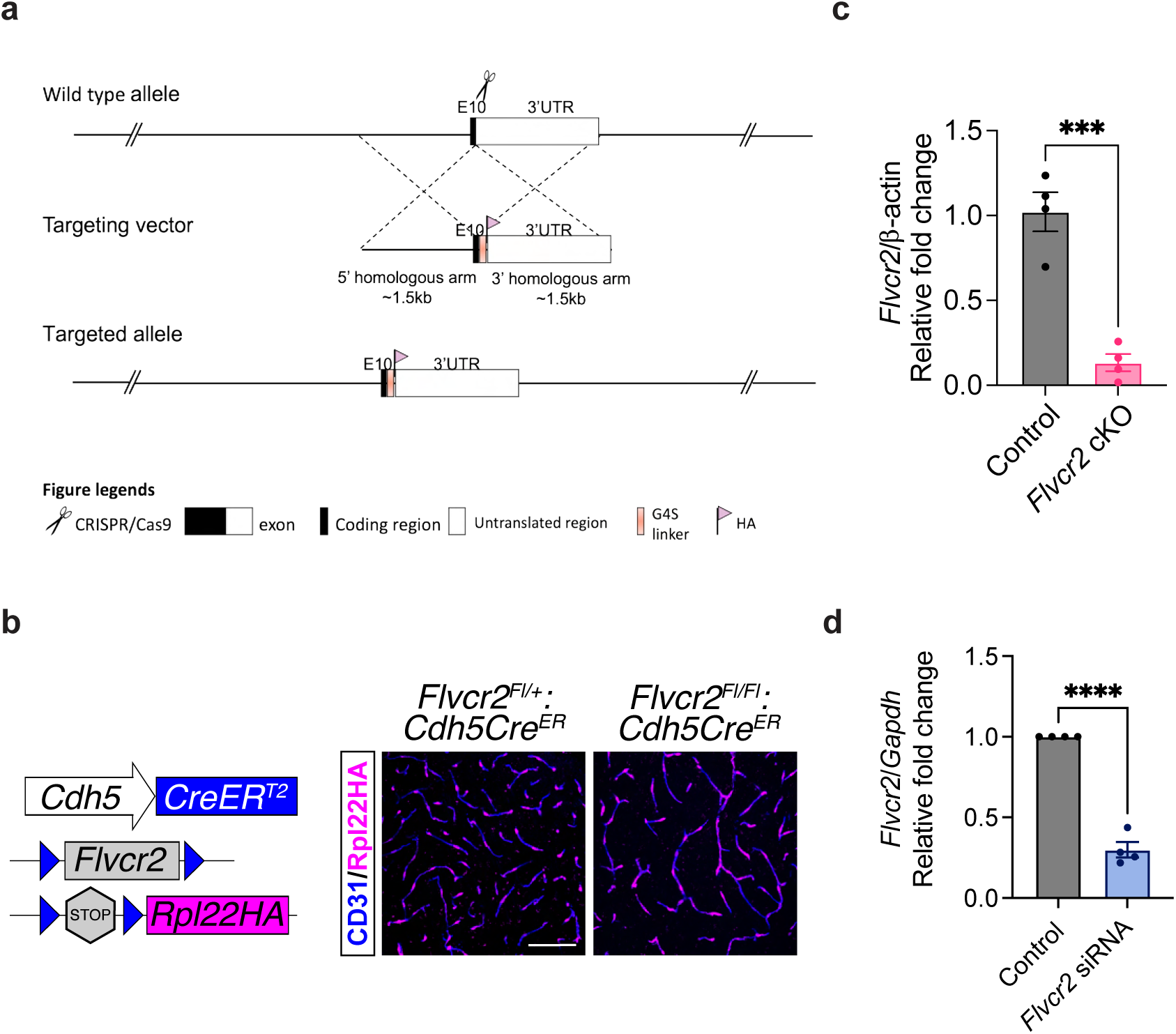
Generation and validation of *Flvcr2* HA-tagging, knock-out, and knock-down. **a,** Targeting strategy to generate *Flvcr2-HA* mice. **b**, Strategy adopted to generate endothelial cell-specific *Flvcr2* conditional knockouts (*Flvcr2* cKO). Brains from control (*Flvcr2^Fl/+^:Cdh5Cre^ER^*) and *Flvcr2* cKO (*Flvcr2^Fl/Fl^:Cdh5Cre^ER^*) mice were stained for HA (Rpl22HA recombination reporter; pink) and CD31 (blue) to delineate blood vessels. Scale bar is 100 µm. These data show successful endothelial-cell specific recombination. **c,** qRT-PCR performed with RNA extracted from adult mouse brain showing control (*Flvcr2^Fl/+^:Cdh5Cre^ER^*) vs *Flvcr2* cKO (*Flvcr2^Fl/Fl^:Cdh5Cre^ER^*) deletion of *Flvcr2*. Mouse β-actin expression was used as the internal loading control. Data are mean ± s.e.m (n = 4); *** indicates p < 0.001. **d,** qRT-PCR performed with RNA extracted from HBEC cells transfected with either scr siRNA (control) or *Flvcr2* siRNA (KD) showing decreased expression of *Flvcr2*. Human *Gapdh* expression was used as the internal loading control. Data are mean ± s.e.m. (n = 4); **** indicates p < 0.0001.

**Extended Data Figure 3.**
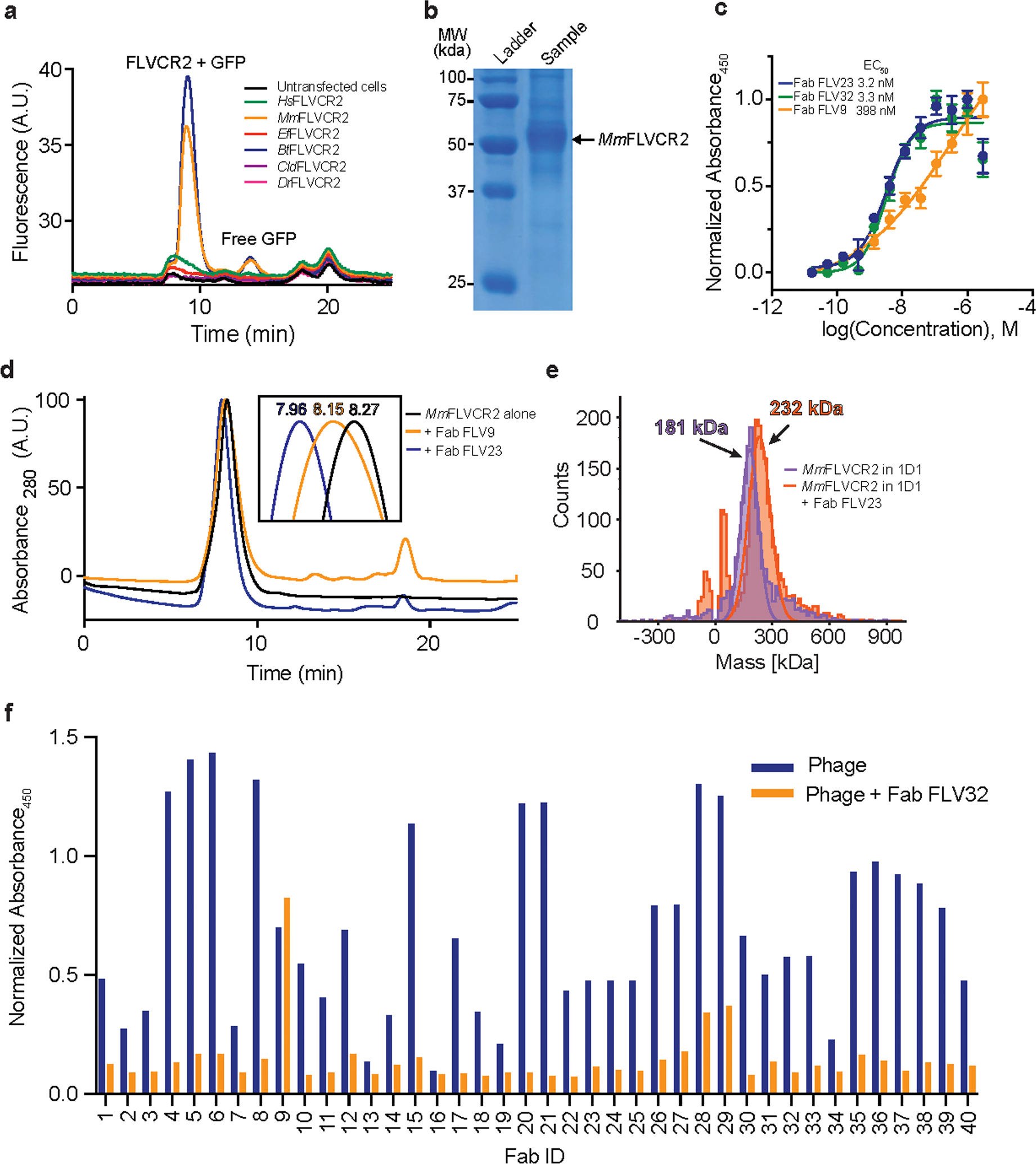
Expression, purification, and Fab complex formation of *Mm*FLVCR2. **a,** Fluorescent size exclusion chromatography elution profiles of six FLVCR2 orthologs fused to green fluorescent protein (GFP). Orthologs screened are from *Homo sapiens* (*Hs*FLVCR2; NCBI – NP_060261.2), *Mus musculus* (*Mm*FLVCR2; NCBI – NP_663422.1), *Eptesicus fuscus* (*Ef*FLVCR2; NCBI – XP_027997147.2), *Canis lupus dingo* (*Cld*FLVCR2; NCBI – XP_025299773.1), *Bos taurus* (*Bt*FLVCR2; NCBI – NP_001179072.1), and *Danio rerio* (*Dr*FLVCR2; NCBI – XP_693589.3). **b,** Representative SDS-PAGE gel of *Mm*FLVCR2 purified in DDM/CHS. **c,** EC_50_ evaluation of select purified Fabs binding to *Mm*FLVCR2 incorporated into MSP1D1 nanodisc. Data points represent the mean ± S.D. (n = 2). **d,** Normalised high-performance liquid chromatography elution profiles of *Mm*FLVCR2 reconstituted into MSP1D1 nanodisc alone (black) and in complex with Fab FLV23 (blue) and Fab FLV9 (orange). Numbers indicate the retention time (mins) of each peak. **e,** Mass photometry analysis of nanodisc-reconstituted *Mm*FLVCR2 purified via size exclusion chromatography in the presence (orange) and absence (purple) of Fab FLV23. The peaks differ by 51 kDa which corresponds to the molecular weight of Fab FLV23. **f,** Single-point phage ELISA for binding of *Mm*FLVCR2-specific Fab-phage in the presence (orange) and absence (blue) of 1 μM purified Fab FLV32, which was used as an epitope masking reagent. Note that FLV32 was confirmed to have the same binding epitope as FLV23 via preliminary cryo-EM experiments (data not shown). Only Fab FLV9 produced the same level of ELISA signal in the presence and absence of the Fab FLV32.

**Extended Data Figure 4.**
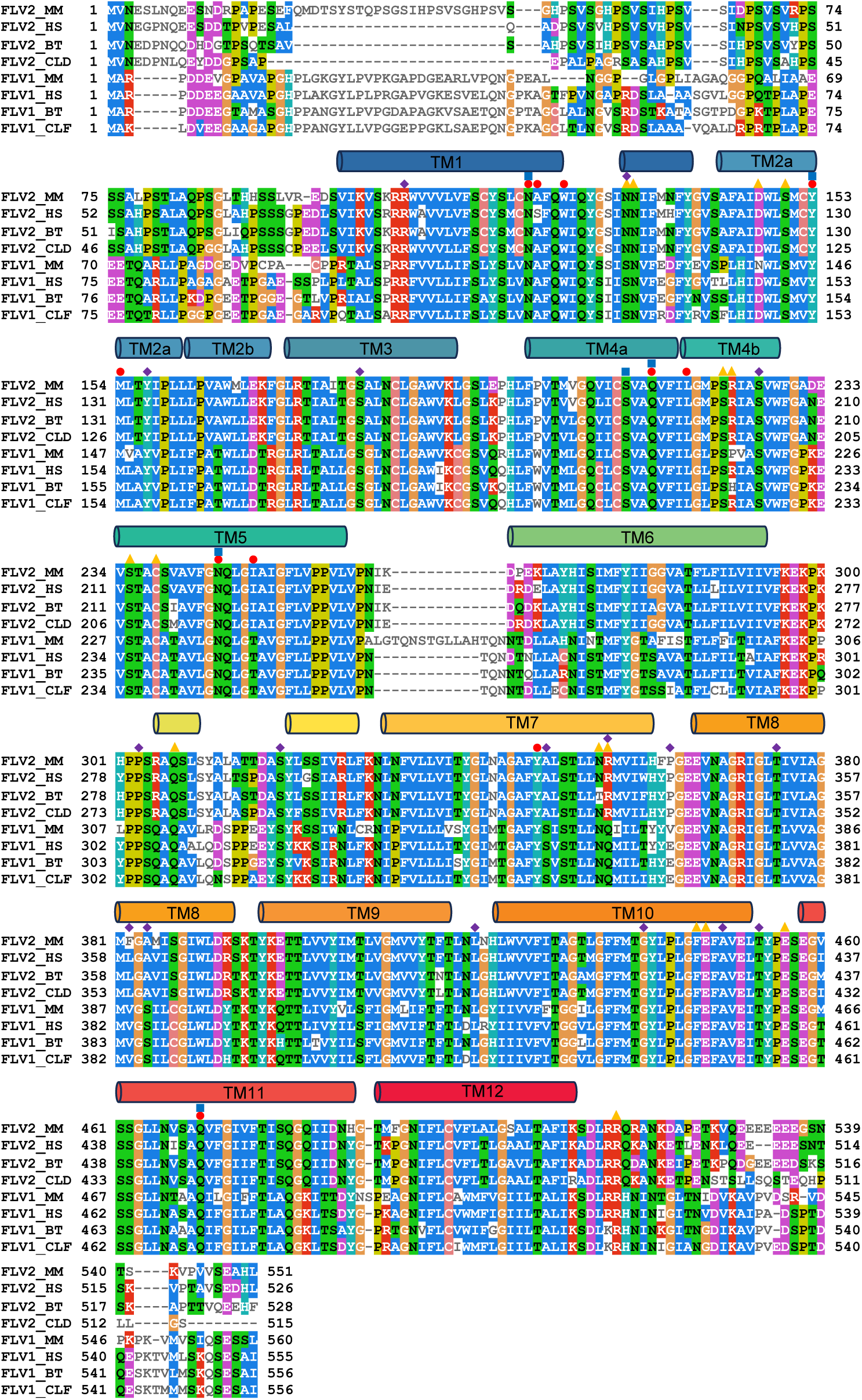
FLVCR2 and FLVCR1 multiple sequence alignment. Four FLVCR2 and FLVCR1 variants were aligned using Clustal Omega^40^ and visualised and coloured using MView, colouring by property colour scheme. The sequences aligned are: *Homo sapiens* (*Hs*FLVCR2; NCBI – NP_060261.2), *Mus musculus* (*Mm*FLVCR2; NCBI – NP_663422.1), *Canis lupus dingo* (*Cld*FLVCR2; NCBI – XP_025299773.1), *Bos taurus* (*Bt*FLVCR2; NCBI – NP_001179072.1), *Homo sapiens* (*Hs*FLVCR1; NCBI – NP_054772.1), *Mus musculus* (*Mm*FLVCR1; NCBI – NP_001074728.1), *Canis lupus familiaris* (*Clf*FLVCR1; NCBI – XP_003639227.1), and *Bos taurus* (*Bt*FLVCR1; NCBI – NP_001192948.1). Secondary structural elements are shown as cylinders, labelled, and coloured as in Fig. 2. Red circles and orange triangles denote choline-coordinating residues and gating residues respectively. Purple diamonds denote residues at which mutations are associated with PVHH. Blue squares represent residues surrounding the substrate-adjacent ion-like cryo-EM densities.

**Extended Data Figure 5.**
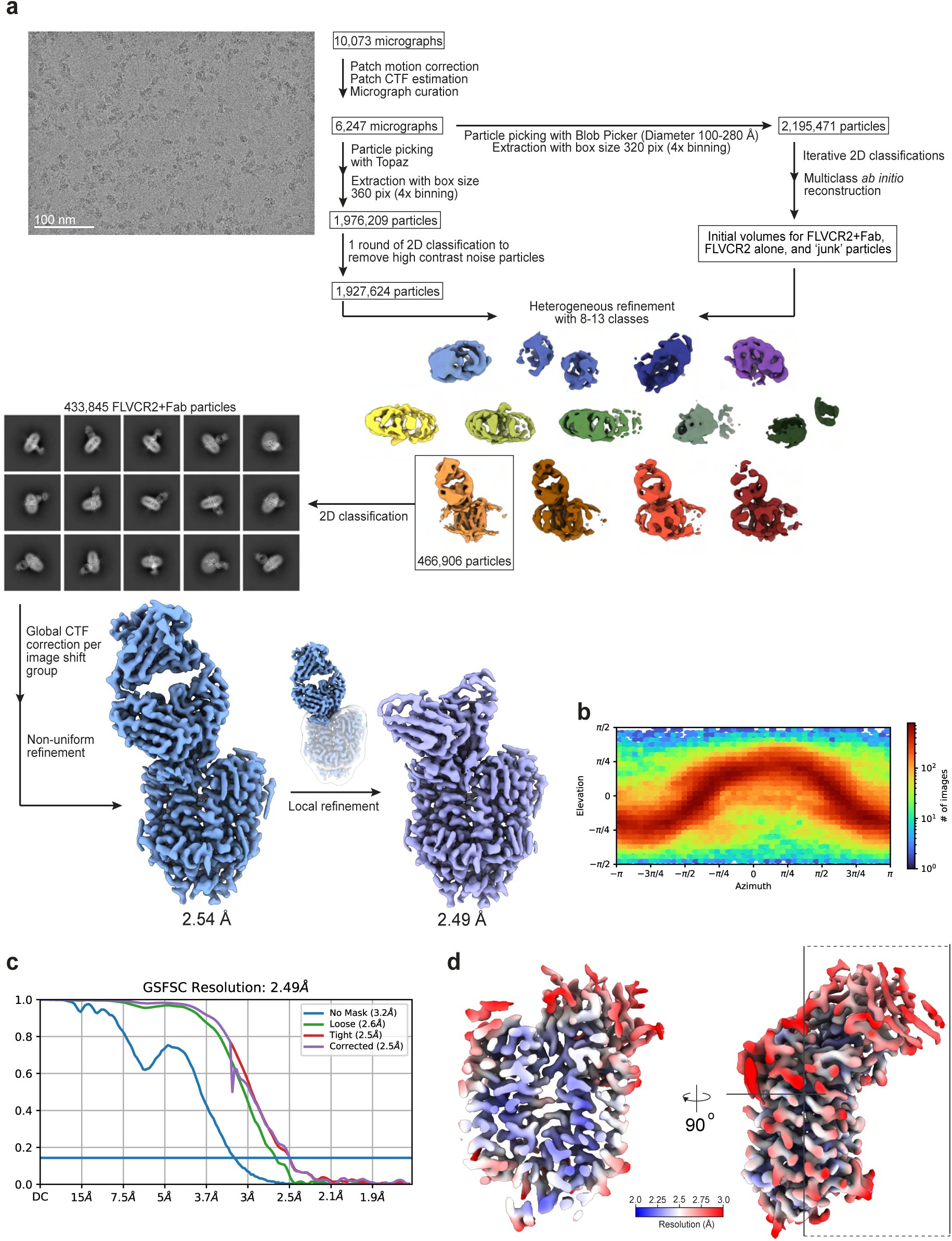
Cryo-EM workflow and analysis of the outward-facing state of *Mm*FLVCR2. **a,** Flow chart outlining cryo-EM image acquisition and processing performed to obtain a structure of detergent-purified *Mm*FLVCR2 in complex with the Fab FLV23. A representative micrograph and 2D class averages are shown. All processing was performed using CryoSPARC v.4.1.2^56^ (see Methods for details). **b,** Euler angle distribution plot of the final three-dimensional reconstruction of the *Mm*FLVCR2-Fab FLV23 complex. **c,** Fourier shell correlation (FSC) curves for the *Mm*FLVCR2-Fab FLV23 complex. **d,** Local resolution map of the *Mm*FLVCR2-Fab FLV23 complex, with an orthogonal view indicating the location of the clipping plane. Density is coloured by resolution from 2.0 (blue) to 3.0 Å (red).

**Extended Data Figure 6.**
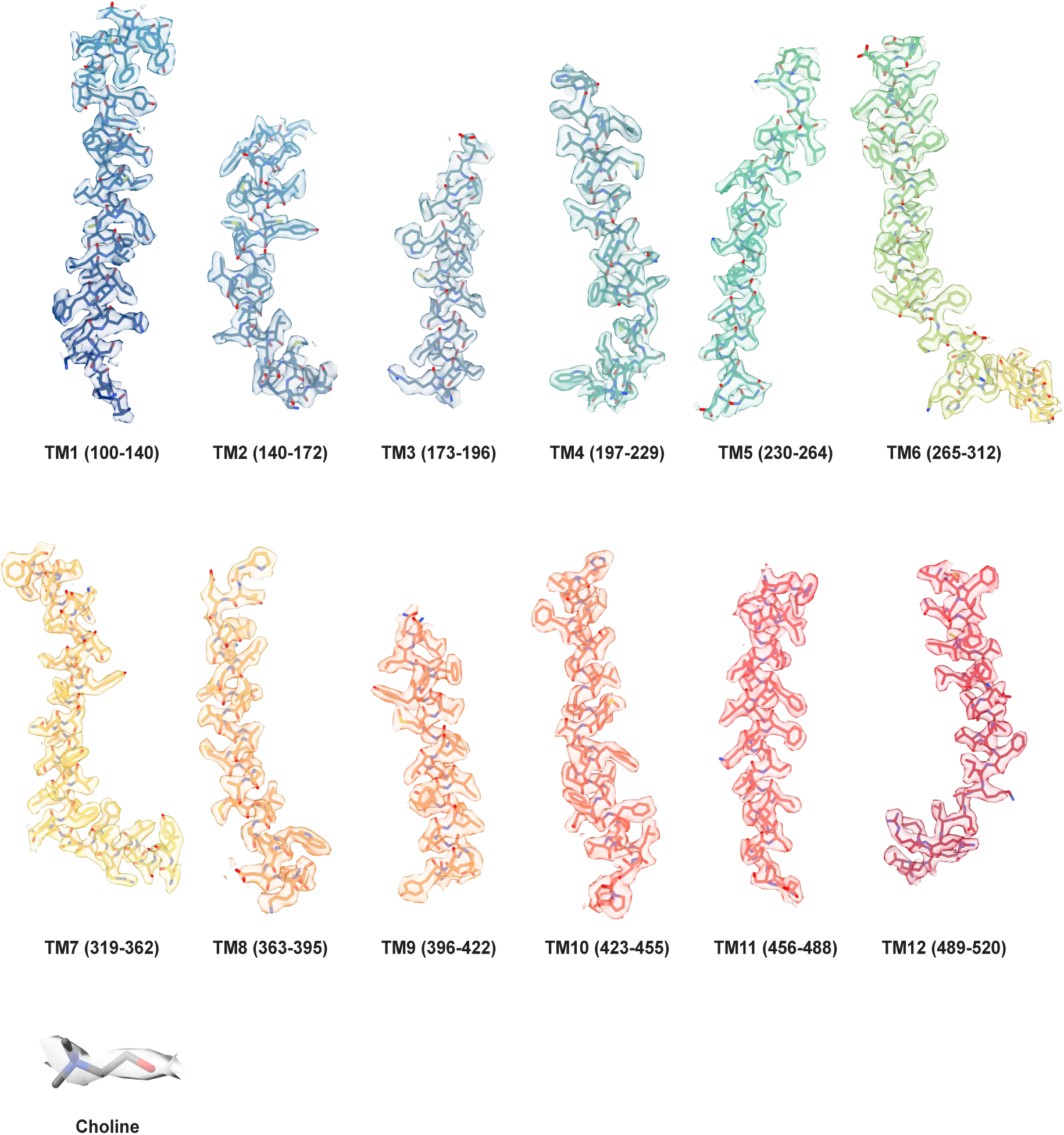
Cryo-EM density fit of the outward-facing FLVCR2 model. Cryo-EM densities (semi-transparent surface) are superimposed on the structural elements of *Mm*FLVCR2 including TM helices 1-12 and bound choline. All elements are shown in stick representation and coloured as in Fig. 2.

**Extended Data Figure 7.**
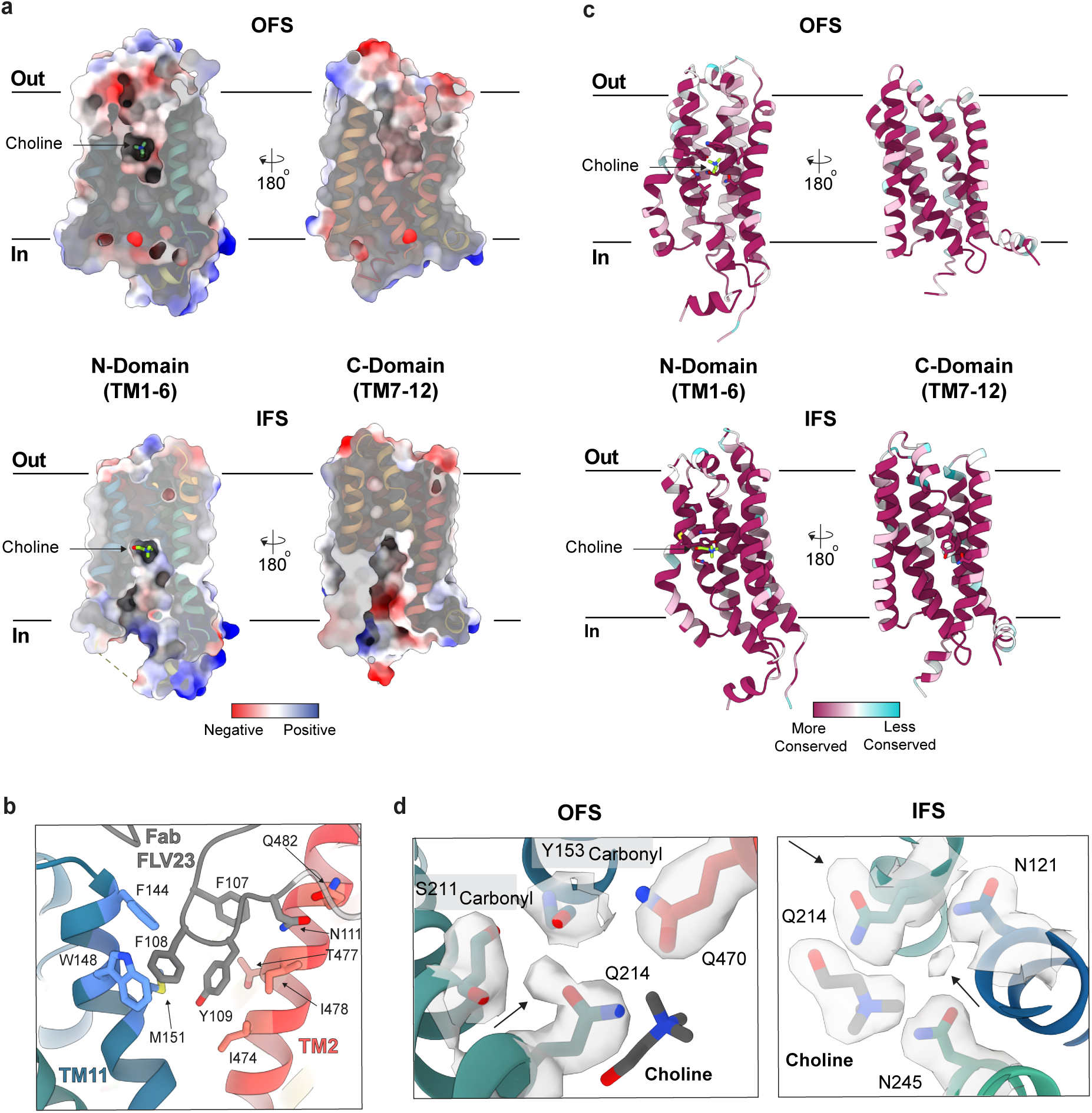
Structural features of FLVCR2. **a,** The N- and C-domains of FLVCR2 as viewed from within the extracellular cavity of the OFS (top panel) and the intracellular cavity of the IFS (bottom panel). For both domains, the electrostatic potential is shown as a surface coloured from red (negative) to blue (positive) with the structure overlayed in ribbon representation and coloured as in Fig. 2. **b,** Interaction site of Fab FLV23 with FLVCR2 in the OFS. TM2, TM11, and Fab FLV23 are coloured red, blue, and grey respectively, with interacting residues highlighted in stick representation. **c,** The N- and C-domains of FLVCR2 viewed in the same manner as in panel **a,** with the protein shown in ribbon representation and coloured by sequence conservation across mammalian FLVCR2 species using a gradient from most (maroon) to least (cyan) conserved. **d,** Ion-like cryo-EM densities observed in close proximity to the choline binding site in the OFS (left) and IFS (right). The protein is coloured as in Fig. 2 with choline shown in grey stick representation and the ion-like cryo-EM densities and cryo-EM density of surrounding residues and choline shown as a grey semi-transparent surface. Positions of the ion-like densities are indicated by the arrows. Note that the ion-like density in the OFS has a higher occupancy than choline.

**Extended Data Figure 8.**
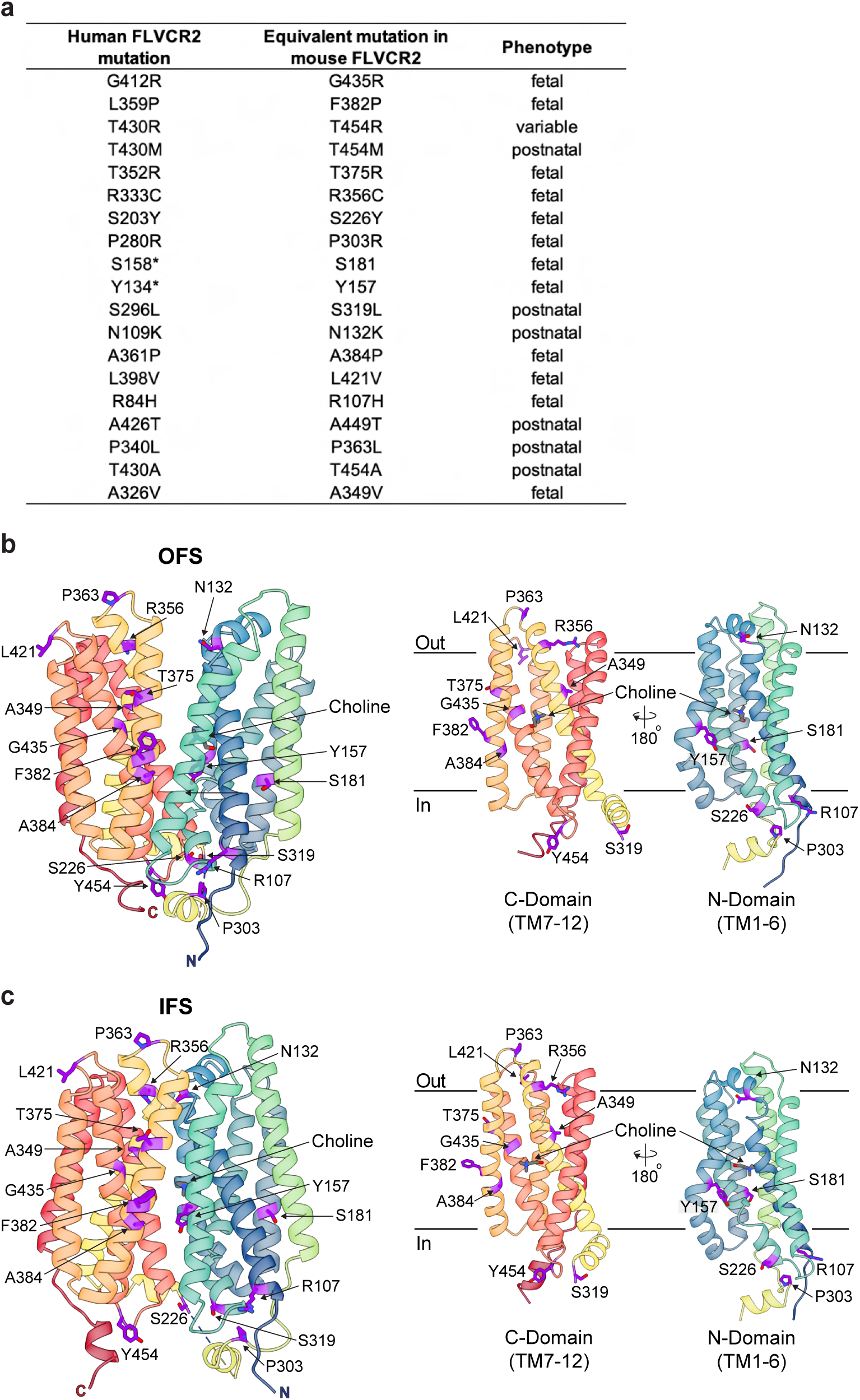
Human mutations in FLVCR2 associated with PVHH. **a,** Known human mutations in FLVCR2 that are associated with PVHH, their equivalent sites in *Mm*FLVCR2, and observed phenotypes associated with these mutations. Postnatal phenotype refers to a mild phenotype, whereas foetal phenotype refers to subjects who died during birth or by compassionate abortion. The position of human mutations in FLVCR2 mapped onto the structure of *Mm*FLVCR2 in the **b,** OFS and **c,** IFS. Left panel shows a front-on view of the protein and the right panel shows the N- and C-domains as viewed from within the central cavity. All views are in the plane of the membrane. The structures are show in ribbon representation and coloured as in Fig. 2, with equivalent sites of human mutations coloured in purple and shown in stick representation.

**Extended Data Figure 9.**
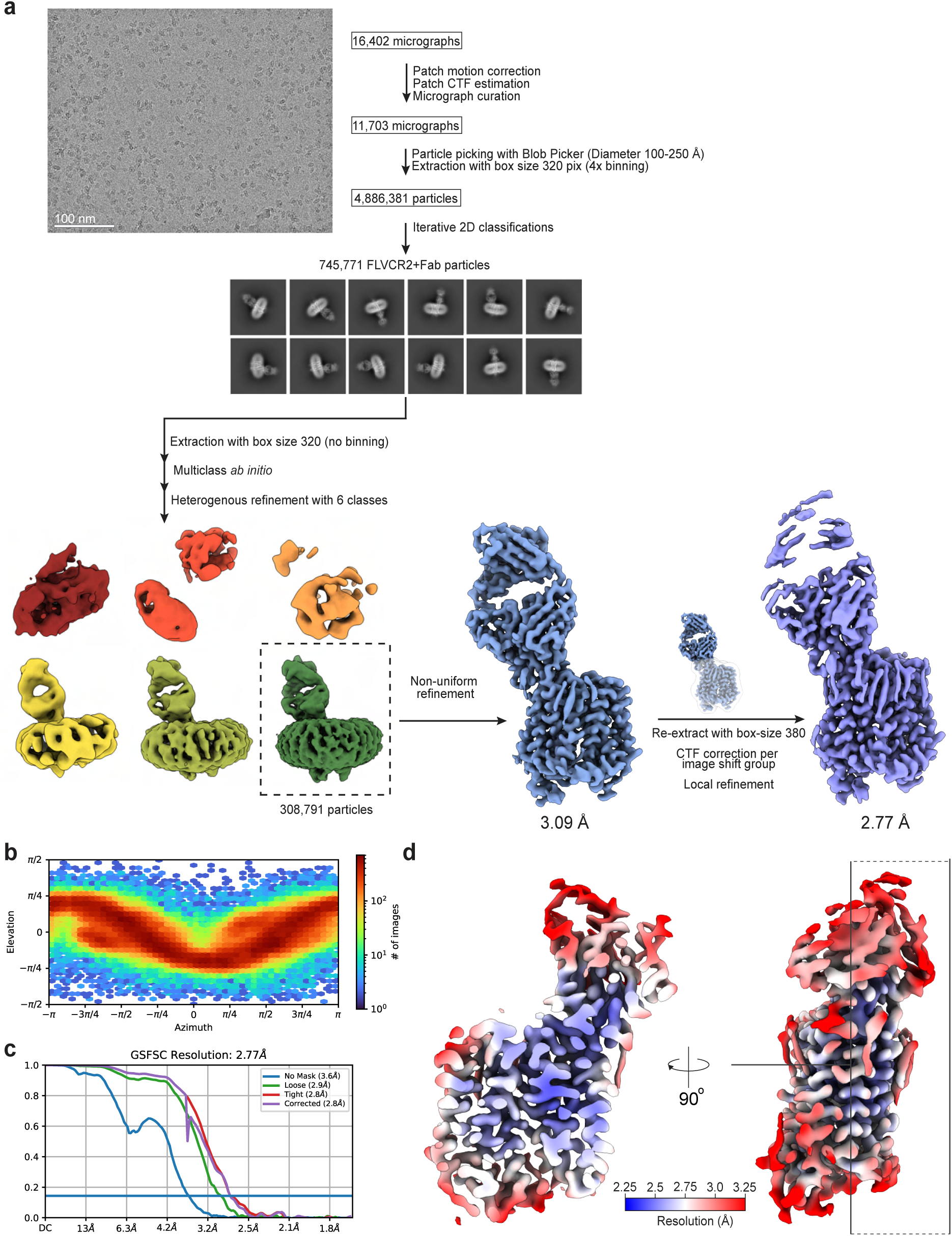
Cryo-EM workflow and analysis of the inward-facing state of *Mm*FLVCR2. **a,** Flow chart outlining cryo-EM image acquisition and processing performed to obtain a structure of detergent-purified *Mm*FLVCR2 in complex with the Fab FLV9. A representative micrograph and 2D class averages are shown. All processing was performed using CryoSPARC v.4.1.2^56^ (see Methods for details). **b,** Euler angle distribution plot of the final three-dimensional reconstruction of the *Mm*FLVCR2-Fab FLV9 complex. **c,** Fourier shell correlation (FSC) curves for the *Mm*FLVCR2-Fab FLV9 complex. **d,** Local resolution map of the *Mm*FLVCR2-Fab FLV9 complex, with an orthogonal view indicating the location of the clipping plane. Density is coloured by resolution from 2.25 (blue) to 3.25 Å (red).

**Extended Data Figure 10.**
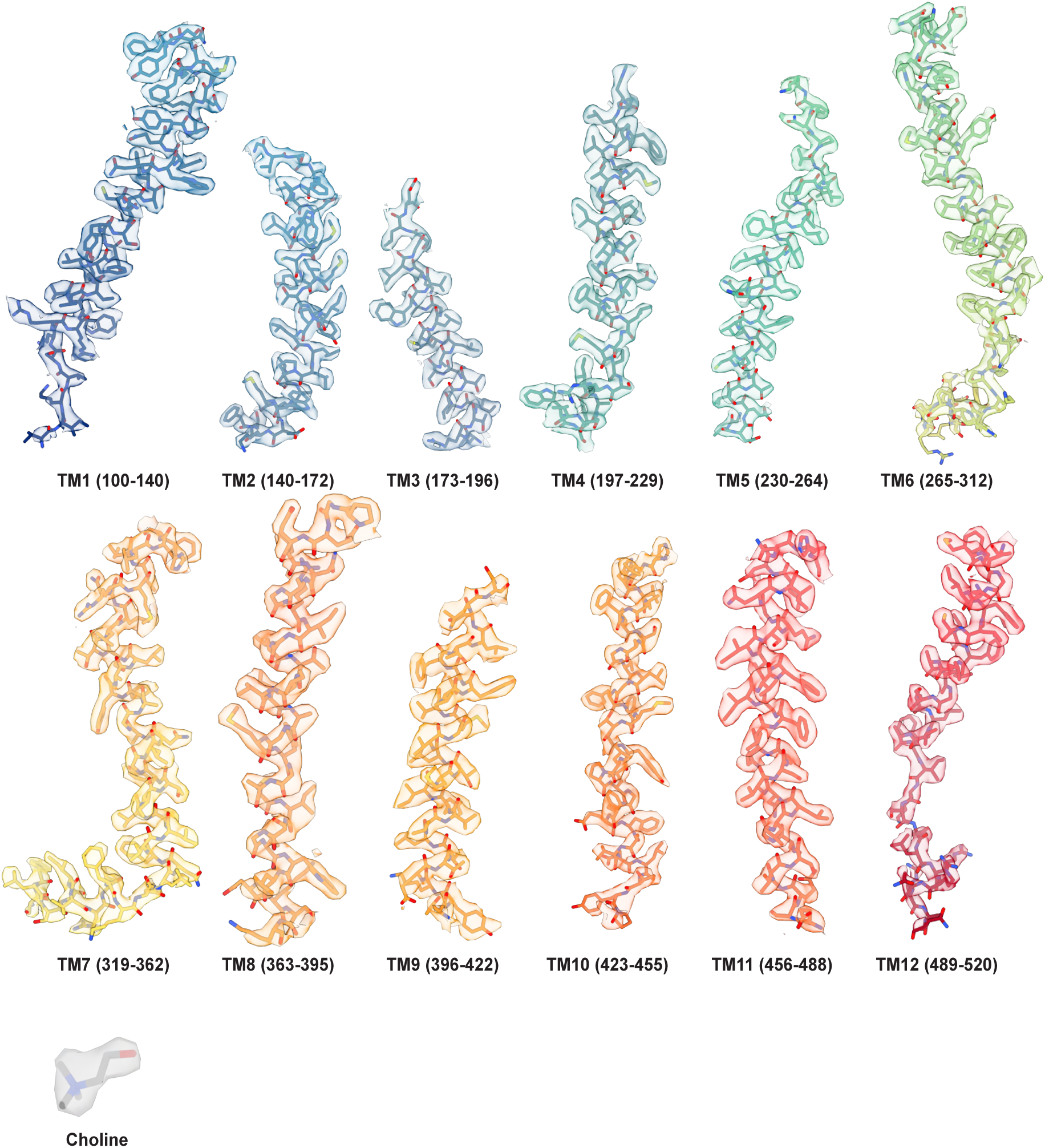
Cryo-EM density fit of the inward-facing *Mm*FLVCR2 model. Cryo-EM densities (semi-transparent surface) are superimposed on structural elements of *Mm*FLVCR2 including TM helices 1-12 and bound choline. All elements are shown in stick representation and coloured as in Fig. 2.

